# Ibuprofen impairs fracture healing by disrupting neutrophil-mediated tissue repair

**DOI:** 10.64898/2026.09.05.749610

**Authors:** Jérémie Zappia, Lucy M. McGowan, Rebecca M. Chatwin, Rabia Sevil, Bianca H. Fernandes, Benjamin T. Gillard, Joanna J. Moss, Christopher P. Jones, Stephen Cross, Nilabhra R. Das, John P. Kemp, Borko Amulic, Chrissy L. Hammond

**Affiliations:** School of Biochemistry and Biomedical Sciences, Biomedical Sciences Building, University of Bristol, BS8 1TD, United Kingdom; Translational Health Sciences, Bristol Medical School, Dorothy Hodgkin Building University of Bristol, BS8 1TD, United Kingdom; Interface Analysis Centre, School of Physics, H. H. Wills Physics Laboratory University of Bristol, Bristol, BS8 1TL, UK; Wolfson Bioimaging Facility, Biomedical Sciences Building, University of Bristol, BS8 1TD, United Kingdom; Mater Research Institute, The University of Queensland, Translational Research Institute, Woolloongabba, Queensland, Australia

## Abstract

Neutrophils are among the first immune cells recruited to fractures, yet their contribution to bone repair remains poorly understood. Anti-inflammatory use has been linked to increased risk of non-union fractures, while prolonged neutrophil activity or failed clearance of neutrophil-derived DNA in the form of neutrophil extracellular traps (NETs), can also impair fracture healing. Using dynamic live imaging and cellular perturbation in adult zebrafish, we show that neutrophils play a key role in shaping the early fracture microenvironment. Neutrophils rapidly accumulated at fractures where they released NETs and exhibited the activation of gene programmes associated with matrix remodelling. Compromising neutrophil recruitment, via ibuprofen treatment or orthogonal genetic and pharmacological perturbations, dramatically increased incidence of fracture non-union and lead to altered callus architecture through delay of osteoblast differentiation and impaired mineralisation. Mechanistically, neutrophils actively interacted with key structural proteins, including laminin, fibronectin, and collagen I, through uptake, recycling and active de novo secretion pathways. Together, our findings reveal that neutrophils establish a provisional emergency fracture matrix that templates subsequent bone repair, providing a mechanistic link between early non-steroidal anti-inflammatory drug (NSAID) exposure and impaired skeletal healing.

## Introduction

Bone fractures represent a major healthcare burden with over 178 million fractures per annum; predicted to rise yet further over the coming years^1^. In Western countries, osteoporotic and fragility fractures account for more hospital days among older adults than diabetes, cardiovascular disease, and cancer combined^2^.

The inflammatory response is among the earliest and most critical determinants of tissue repair. Acute inflammation protects injured tissues from infection and orchestrates the recruitment of cells required for tissue repair. However, excessive or prolonged inflammatory signalling can compromise repair outcomes^3^ This creates a clinical dilemma: although suppression of inflammation is a common therapeutic strategy after musculoskeletal injury, accumulating evidence suggests that non-steroidal anti-inflammatory drug (NSAID) use may adversely affect fracture healing and contribute to non-union repair^4–6^

The early acute pro-inflammatory environment is predominantly driven by neutrophils, which are among the earliest immune cells to infiltrate injured tissue. While historically neutrophils have been solely viewed as mediators of acute inflammation and chronic immunopathology, increasing evidence indicates neutrophils can also promote tissue repair^7,8^, and disruption of neutrophil function has been associated with impaired skeletal repair^9^. Early NSAID administration could unintentionally suppress the neutrophil-driven repair in the earliest phases of repair when establishment of a pro-reparative niche is beneficial.

How exactly early immune cells influence the subsequent behaviour of bone secreting cells, or osteoblasts, during bone regenerative processes remains unclear. Recent evidence suggests neutrophils participate in tissue repair by remodelling the injury microenvironment^10^. Successful regeneration requires more than the delivery of signalling cues; it requires that cells respond to these cues to establish a microenvironment which is permissive for repair, which in matrix rich tissues, such as bone, will involve remodelling of the local matrix. The extracellular matrix (ECM) is no longer viewed as a passive scaffold but as a dynamic regulator of cell migration and behaviours, which contribute to the generation of an environment conducive to regeneration^11^. Innate immune cells play a role in establishing the regenerative environment, macrophages can both process local matrix and interact with secretory cells to influence matrix deposition; in fracture recent spatial omics findings suggest that crosstalk between macrophages and osteoblasts could coordinate repair^12^. Further recent data has shown that neutrophils release NETs after fracture in mice and humans, and that blocking NET formation slows mineralisation, but that sustained NET presence leads to poor outcomes, suggesting that timing of neutrophil activity is a critical factor^13^. Neutrophils in the hematoma, the clot which forms after fracture, have been shown to contain matrix proteins, including fibronectin, which is postulated to help provide an emergency scaffold^14^. Although neutrophils are increasingly recognised as important regulators of tissue repair and ECM remodelling^15^, whether and how they directly contribute to the establishment of a regenerative microenvironment remains unknown.

Here, we identify neutrophils as previously unrecognised architects of the fracture microenvironment. NSAID treatment depletes neutrophils at the injury site, resulting in defective bone repair marked by delayed osteoblast differentiation and mineralisation. We show that neutrophils construct an early regenerative scaffold through extracellular chromatin released via neutrophil extracellular traps (NETs) that bridges the fracture gap and organises extracellular matrix proteins. Beyond matrix organisation, neutrophils actively synthesise and internalise matrix components, revealing an unexpected role in matrix turnover and remodelling. Collectively, our findings establish neutrophils as indispensable early coordinators of the regenerative niche that initiate successful bone repair.

## Results

### Early ibuprofen treatment attenuates injury-induced neutrophil recruitment and associated transcriptional diversification

We used the caudal fin fracture model in adult zebrafish to establish timing of neutrophil recruitment to a fracture injury. In line with previous work showing *lyz+* myeloid cells are recruited to induced caudal fin fractures within the first hours of injury^16^we established timing of neutrophil recruitment using the *mpx:GFP* transgenic reporter, in which cytosolic GFP is produced by cells expressing neutrophil myeloperoxidase^17^. Fractures were induced in the caudal fins of adult *mpx:GFP* zebrafish, which resulted in robust neutrophil accumulation at the fracture site from 2 hpi (hours post injury), peaking at 4 and 8 hpi, and waning by 24 hpi, with numbers restored to baseline by 48 hpi (Supplementary Figure 1).

To determine the consequences of NSAIDs on neutrophils we treated fish with ibuprofen (10μM), corresponding to physiologically relevant dosages found in human plasma^18^ for 24 hours prior to fracture induction and then continuously thereafter, while simultaneously longitudinally tracking the neutrophil response within the same injuries. Ibuprofen significantly reduced neutrophil recruitment to the fracture site at 8 hpi (Fig 1A, B). Despite the reduced numbers, those neutrophils which did arrive produced reactive oxygen species (ROS) (Supplementary Fig 2). Non-union can easily be quantified in zebrafish, as it leads to the loss of the unattached ray bone, followed by subsequent regeneration. Strikingly, we observed an increase in frequency of non-union events following ibuprofen treatment (p = 0.0263), such that 44.44% of treated fish did not achieve union compared to 18% of control fish (Fig 1C, D). Non-union events were characterised by reduced osteoblast area (marked by *sp7/osterix*, a master regulator of the osteoblast lineage) in ibuprofen treated fish, compared to DMSO treatment (Fig 1E), suggesting that ibuprofen blunts osteoblast differentiation following injury.

**Figure 1:**
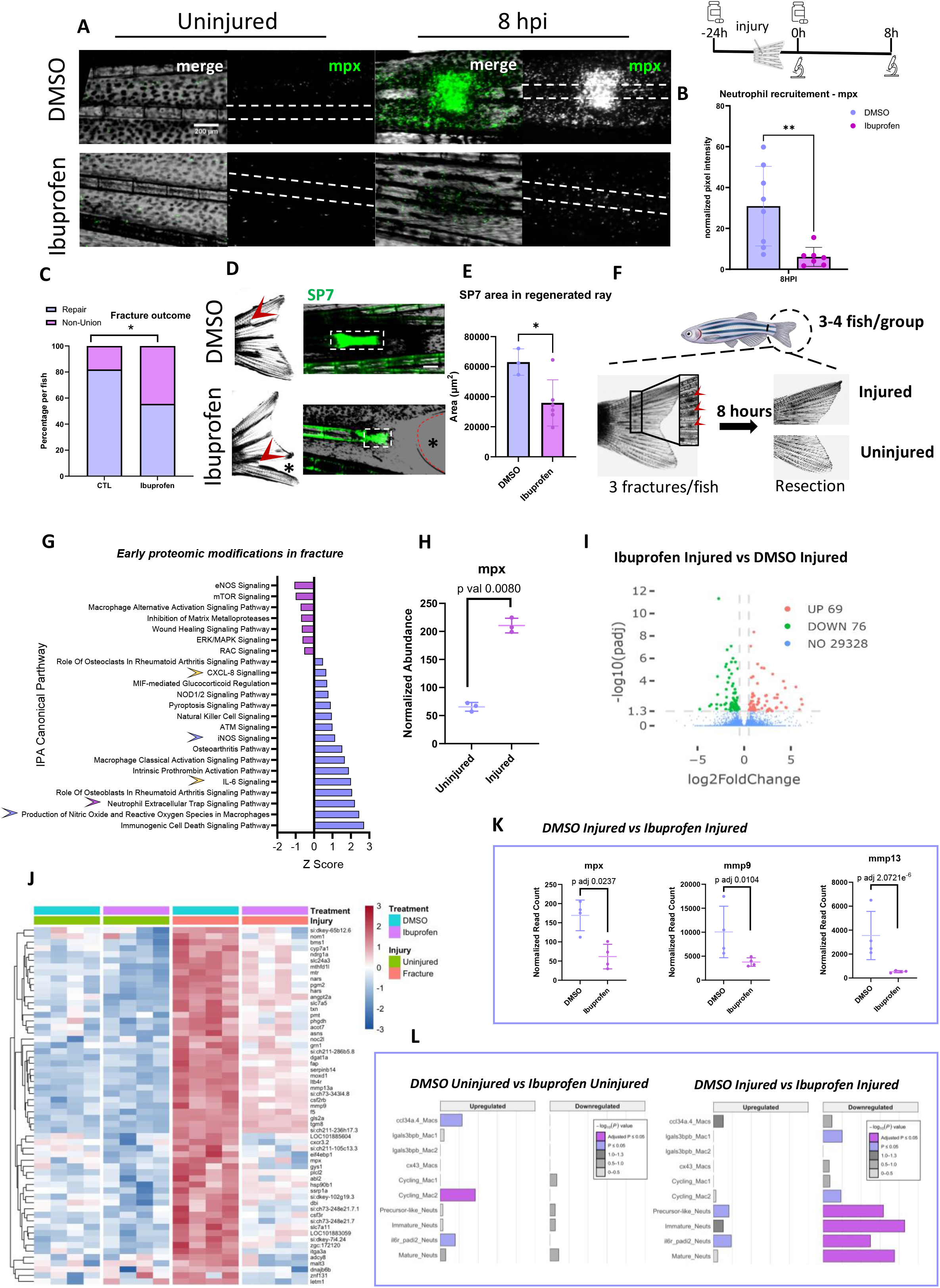
A neutrophil signature is enriched during the early response to bone injury, whereas ibuprofen treatment (10 μM) reduces neutrophil recruitment to the injury site and diminishes the enrichment of genes associated with heterogeneous neutrophil cell states. **A:** Representative stereomicroscope image of uninjured and injured ray at 8 hpi in *Tg(mpx:GFP)* zebrafish treated with Ibuprofen compared to control (DMSO). Scale bar = 200 μm. **B:** The normalized pixel intensity for *mpx:GFP* is plotted as a bar plot with SD. Statistical analysis was performed with a Welch’s test, n=7-8 per group. ** = P < 0.01. **C:** Percentage of the incidence of non-union fracture following Ibuprofen treatment, per individual. Statistical analysis was performed with a Chi-square, n = 50 for the control group and n = 18 for the ibuprofen group. **D:** Representative stereomicroscope image of non-union fracture in *sp7:nlsGFP* DMSO control and ibuprofen treated fish. Red arrowhead indicates the non-union fracture and the (*) shows impaired wound healing. Scale bar = 200 μm. **E:** The regenerated bone area (μm^2^), indicated by the box with dash line in (D), was measured at 3 dpi and plotted as a bar plot with SD, n=3-6 per group. Statistical analysis was performed with a Welch’s test, * = P < 0.05. **F:** Schematic representation of the sample collected for transcriptomic and proteomic. On the dorsal half of the caudal fin, 3 fractures were induced on 3 different bony ray (red arrowheads). The ventral portion of fins were left uninjured as internal controls. Uninjured and injured samples were harvested at 8 hpi to allow neutrophil recruitment. For transcriptomic analysis, 4 groups of 3 fish were pooled together and for proteomic analysis, 3 groups of 3 fish were pooled together. **G:** IPA analysis from proteomic of injured vs uninjured fish at 8 hpi. Yellow arrowheads show inflammatory signature. Blue arrowheads show ROS related signature. Purple arrowhead shows NETosis signature. The normalized abundance of mpx is plotted with SD. **H:** Normalized abundance of *mpx* plotted with SD from proteomic comparison of injured vs uninjured fins. **I:** Vulcano plot of ibuprofen vs DMSO in injury (p-adj < = 0.05 and |log_2_foldchange|>= 0.5. **J:** Heatmap analysis of ibuprofen vs DMSO groups, including injured and uninjured control (p-adj < = 0.05 and |log_2_foldchange|>= 0.5). **K:** Comparison of neutrophil related DEG (*mpx*, *mmp9*, and *mmp13*) in ibuprofen vs DMSO in injury. Normalized read count are plotted with SD. **L :** Myeloid cluster enrichment of genes differentially expressed following ibuprofen treatment. Hypergeometric enrichment of differentially expressed genes from ibuprofen vs DMSO comparisons in uninjured (left) and injured (right) samples within cluster-specific marker gene sets from the published zebrafish burn-injury scRNA-seq dataset [13]. Upregulated and downregulated panels represent bulk RNA-sequencing genes classified by their log2FC in the ibuprofen vs DMSO comparison: upregulated genes were expressed at higher levels in ibuprofen treated samples (log2FC ≥ 0.5), whereas downregulated genes were expressed at lower levels in ibuprofen-treated samples, and therefore at higher levels in DMSO controls (log2FC ≤ −0.5). Bar length represents −log10 of the unadjusted hypergeometric P value. Purple indicates Bonferroni-adjusted P ≤ 0.05, light blue indicates nominal P ≤ 0.05, and grey shades indicate nonsignificant −log10(P) ranges.

All non-union events occurred within the first 8 hours, suggesting the first hours after injury are a key determinate of successful fracture repair outcomes (Supplementary Figure 1D). To capture the gene and protein changes during the early injury response, we performed bulk-omics analyses at 8 hpi, comparing injured tissue with uninjured distal tissue from the same animals to reduce interindividual variation (Figure 1F, with comparison between distal tissues and injury naive fish in Supplemental Figure 3). We observed a transcriptomic signature related to neutrophil infiltration following fracture with strong upregulation of *mpx* (myeloperoxidase) expression, as well as *cxcl8a* (C-X-C motif chemokine ligand 8a), and *mmp9* (matrix metalloproteinase-9), and of downstream targets such as *mmp13* (matrix metalloproteinase-13), which is implicated in ECM remodelling (Supplemental Figure 4A-C, E). Bulk proteomics further validated a neutrophil signature following injury (Mpx, IL-6, CXCL-8 signalling) and revealed ROS production and NETosis as upregulated effector functions occurring during the initial fracture phase (Fig 1G, H). The early fracture phase was accompanied by the upregulation of osteoblast differentiation pathways including *bmpr1bb*, *tgfb1b*, *tgfbr2a* with concomitant downregulation of inhibitors *bambi-a* and *dkk2* (Supplemental Figure 4D). Interestingly, ECM related genes such as *col1a1a* and *col5a3b (*collagen family); *lamb2* (laminin); and *postnb* (periostin) were downregulated (Supplemental Figure 4E). Conversely, genes involved in control of mineral formation such as *enpp1* (ectonucleotide pyrophosphatase/phosphodiesterase 1) and *entpd1* (ectonucleoside triphosphate diphosphohydrolase 1) were upregulated (Supplemental Figure 4F).

We observed profound changes to the transcriptome of injured control and ibuprofen treated fish (Fig 1 I, J), while ibuprofen treatment had limited effect on the transcriptome of uninjured fish (Supplemental Fig 5A). Ibuprofen treated injured fish showed a reduction in *mpx, mmp9,* and *mmp13* expression compared to vehicle treated injured fish, (Fig 1K). Upregulation of osteoblast genes *bmpr1bb*, *tgfbr2a, tgfb1b;* and mineralisation genes *enpp1* and *entpd1* was maintained at 8hpi in ibuprofen treated fish while osteoblast differentiation pathways inhibitors *bambi-a* and *dkk2* were still downregulated (Supplemental Figure 5B-D). By contrast, fewer differences in ECM gene expression in ibuprofen treated fins were detected following injury, suggesting ibuprofen leads to selective changes to a specific subset of genes (Supplemental figure 5B, E). Over-representation analysis (ORA) using a hypergeometric test showed that gene programs of several neutrophil cell clusters, defined previously by others using single cell transcriptomics ^19^, were enriched for genes that were downregulated in response to ibuprofen treatment in injured 8 hpi fish (Methods, Fig 1L).

Taken together, these data implicate neutrophils as a candidate for mediating the downstream responses to ibuprofen-responses in the early post fracture response.

### Early ibuprofen administration during fracture healing impairs neutrophil recruitment, compromising bone repair and altering callus formation

To establish the impact of NSAID treatment on bone repair outcomes, we followed osteoblast localisation and differentiation during callus formation, using longitudinal imaging with an *sp7:GFP* transgenic reporter line. Exposure to Ibuprofen delayed osteoblast differentiation within the callus, such that *sp7* expression was significantly lower at 3 and 4 days post-injury (dpi) (Fig 2A, B). Calcein green labelling of newly deposited bone showed that delayed osteoblast differentiation was associated with reduced mineralization at 10dpi (Fig 2C, D). High resolution computed tomography of fracture callus 7dpi showed that differences in osteoblast and calcein green were predictive of altered bone morphology and porosity during repair (Fig 2E).

**Figure 2:**
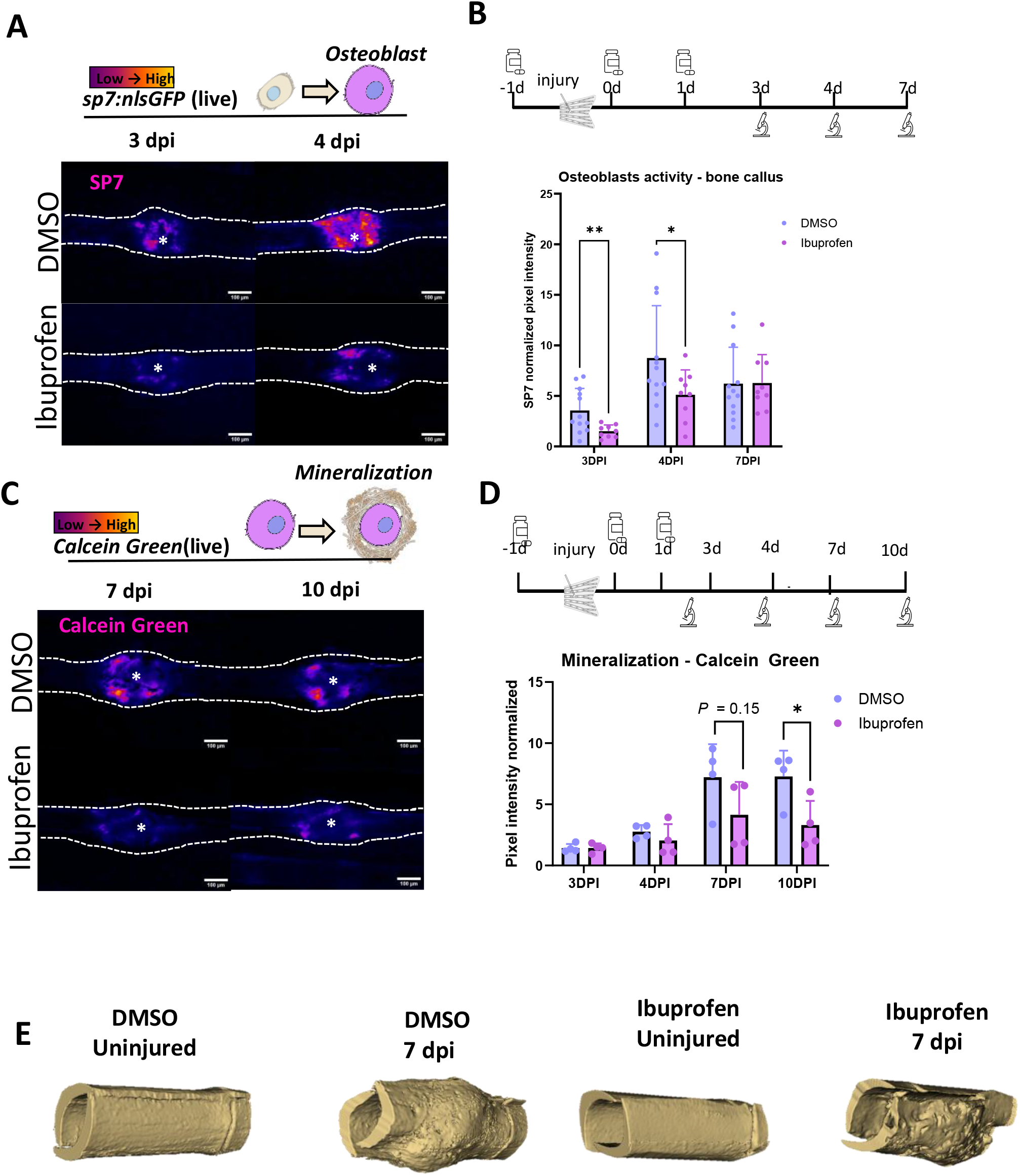
Ibuprofen treatment (10μM) negatively impacts bone repair outcomes. **A:** Representative stereomicroscope images of the intensity of expression of the osteoblast marker *sp7* at 3 and 4 dpi comparing DMSO control and ibuprofen treated fish. The (*) marks the centre of the injury. Scale bar = 200 μm. **B:** Fish were longitudinally followed from 0 to 7 dpi for measuring *sp7* expression. The normalised *sp7* pixel intensity is displayed as a bar plot, n=9-12 per group. Statistical analysis was performed with a two-way Anova, ** = P < 0.01, * = P < 0.05. **C:** Representative stereomicroscope images of the intensity of calcein green staining to assess neomineralization in DMSO control and ibuprofen treated fish. The (*) marks the centre of the injury. Scale bar = 200 μm. **D:** Fish were longitudinally followed from 0 to 10 dpi for measuring calcein green intensity. The normalised pixel intensity is displayed as a bar plot, n=4 per group. Statistical analysis was performed with a two-way Anova, ** = P < 0.01, * = P < 0.05. **E:** Volume rendering of high resolution nanoCT of uninjured bone and bone calluses at 7 dpi.

To confirm that changes to neutrophil behaviour underpinned impaired repair, we used two orthogonal perturbations to reduce neutrophil numbers at the injury site: pharmacological inhibition and genetic perturbation. Treatment with Cxcr1/2 inhibitor SB225002, which inhibits neutrophil chemotaxis^20,21^, significantly reduced neutrophil abundance at the fracture site at 8 hpi (Supplemental Fig 6A, B). It also led to delayed osteoblast differentiation at the fracture site (Supplemental Fig 6C, D), and increased incidence of non-union fracture to 41.66% (Supplemental Fig 6E).

As a further confirmation, neutrophils were depleted prior to injury via ablation in the *Tg(lyz:NTR-mCherry)* line ^22,23^. This reduced *lyz+* neutrophil recruitment to the fracture site by 86.66 % and *mpx+* neutrophils by 91.98% at 8 hpi (Supplemental Fig 7A-C) and led to altered callus formation reminiscent of that seen following ibuprofen treatment (Supplemental Fig 7D, E).

Collectively, our three orthogonal perturbations link reduced neutrophil recruitment during the early post-injury window with later defects in osteoblast differentiation and callus architecture.

### Neutrophils act as architects of the emergency fracture matrix through NET release and matrix management

In injury, neutrophils engage various effector mechanisms, such as the extrusion of NETs, consisting of extracellular chromatin associated with pro-inflammatory and antimicrobial proteins, including myeloperoxidase^24^. Excessive NETosis can contribute to persistent inflammation^25,26^; and more recently, prolonged presence of NETs has been implicated in increased incidence of non-union fracture^27^. Since proteomics revealed a signature of NETosis in fracture at 8 hpi (Fig 1G), we used the myeloid-specific fluorescent histone reporter line *lyz:h2a-mCherry* zebrafish^28^ to follow NETosis *in vivo*. We captured neutrophils extruding their chromatin in real-time at the injury site, (Fig 3A, B, Supplementary video 1A, B). Further evidence of NETosis at the injury site was collected through electron microscopy, identifying enucleated neutrophils, (suggesting they have extruded their nuclear DNA) located near the fracture (Supplemental Fig 8, Supplemental video 2).

**Figure 3.**
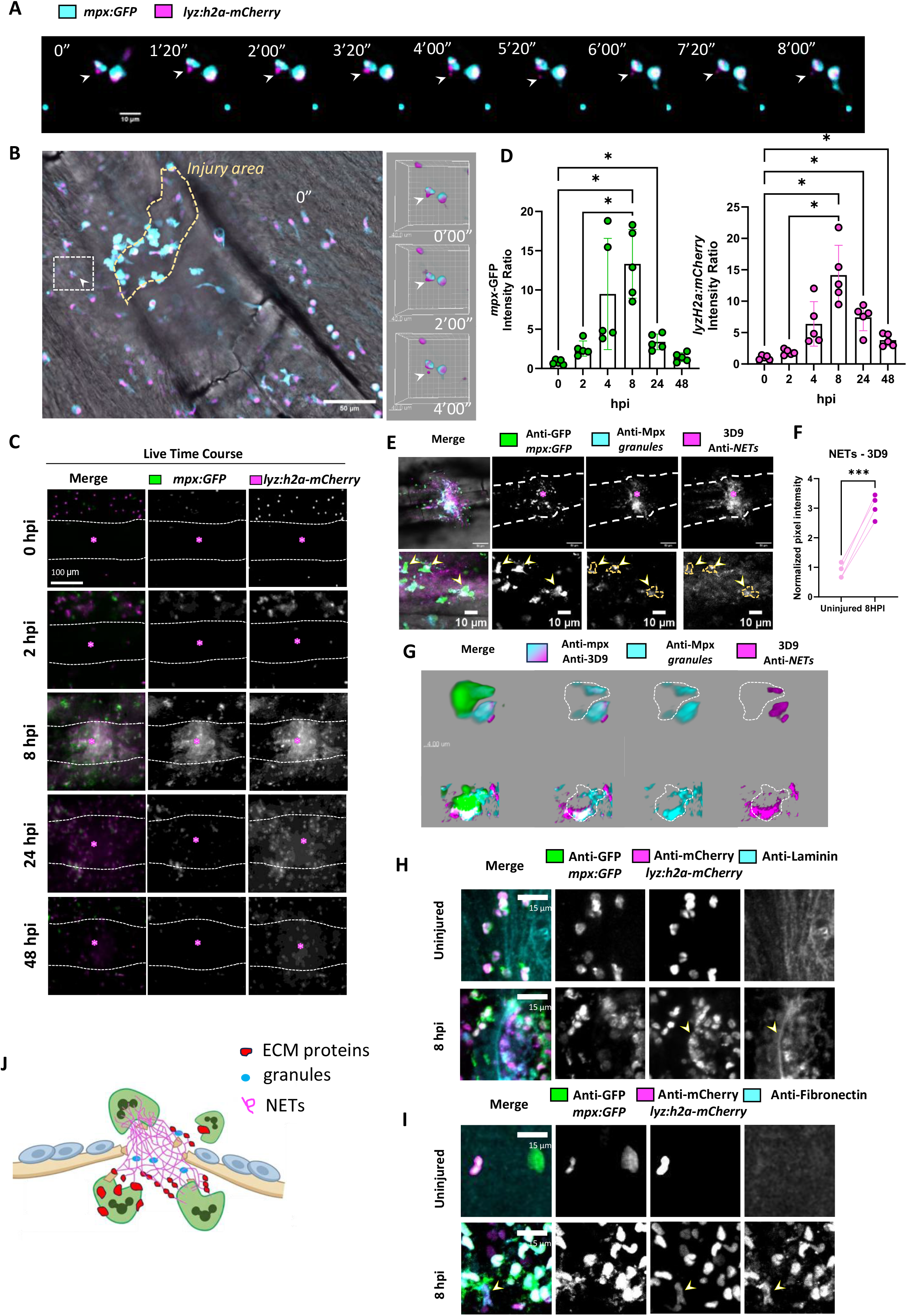
Neutrophils initiate NETosis to remodel the injury environment. **A:** Time lapse isolated from spinning disk imaging (Supplementary video 1) showing *in vivo* NETosis. Fracture was performed in *mpx:GFP; lyz:h2a-mCherry* fish. The caudal fin was resected and imaged every 1’20” approximatively from 4.5 hpi during 2 hours in culture media at 28C. The time lapse shows neutrophils extruding their chromatin *in vivo* (white arrowhead) at proximity to the fracture site. The captured event started approximatively at 5.5 hpi (timestamped 0”). A single z slice was selected to display the timelapse between 0” and 8’. Scale bar = 10 μm **B:** (Left) Image of the fracture with neutrophils recruitment from which the neutrophil performing NETosis was isolated (indicated cell in dotted square). Scale bar = 50 μm. (Right) Imaris 3D volume rendering shows the chromatin coming from within the neutrophils and extruded outside the extracellular space by 4’ (white arrowhead). Scale bar = 40 μm. **C:** Longitudinal observation of *mpx:GFP; lyz:h2a-mCherry* between 0 and 48 hpi to visualise neutrophils and neutrophil-derived chromatin respectively. Dotted line = bone, (*) = centre of fracture. Scale bar = 100 μm. **D:** Quantification of intensity ratios for GFP and mCherry between 0-48 hpi, n =5. Statistical analysis was performed with a Friedman test, *** = P < 0.001, * = P < 0.05. **E:** Representative confocal image at 8 hpi of fracture immunostained for GFP (neutrophils), Mpx (intra and extracellular granules) and NETs (3D9 antibody). Arrowhead indicates neutrophils colocalizing for Mpx and 3D9 signals. Dotted line = bone, (*) = centre of fracture, scale bar = 50 μm (top) and 10 μm (bottom). **F:** The normalized 3D9 pixel intensity was measured at the fracture site and compared to the uninjured bony ray from an internal control fish. Statistical analysis was performed using a Paired t-test, n=4, *** = P < 0.001. **G:** Volume rendering was performed with Imaris, showing the colocalization of mpx granules and NETs secreted by neutrophils. Scale bar = 4 μm. **H-I:** Representative confocal image of uninjured bone and fracture at 8 hpi immunostained for GFP (neutrophils), mCherry (neutrophil-derived chromatin), and laminin (**H**) and fibronectin (**I**). Dense nuclear chromatin and cytosolic GFP can be observed within neutrophils in uninjured tissue. At 8 hpi, large extracellular diffuse regions of neutrophil cytosolic GFP surrounding decondensed neutrophil chromatin indicate NET release (arrowheads). These NETs associate laminin positive regions (**H**) and fibronectin positive regions (**I**). Scale bar = 15 μm **J:** Illustration representing neutrophils under NETosis recruited toward the fracture. Neutrophils secrete NETs to bridge the injury and use it as a scaffold to reorganize the local ECM proteins.

To further establish the presence of NETs we quantified *mpx:GFP; lyz:h2a-mCherry*, colocalization longitudinally in the injury site. Both *mpx* and *lyz* exhibit progressive diffusion, extending beyond cellular boundaries, with extracellular filaments characteristic of NET release (Fig 3C). NET-associated DNA at the injury site perdures longer than the cells, DNA is visible at the fracture site at 24 hpi when neutrophils are less present (Fig 3C, D). To definitively establish these structures as NETs, we co-stained fractures at 8 hpi for GFP to detect neutrophils, the granule component Mpx and 3D9, an antibody which recognises a cleaved fragment of Histone H3 occurring exclusively in NETs^29^, indeed 3D9 and Mpx colocalise at 8hpi (Fig 3E-G, Supplementary video 3).

NET-associated proteases have been previously shown to remodel the matrix proteins laminin and fibronectin^30,31^, therefore, we stained fractures from the transgenic fish with anti-laminin and anti-fibronectin antibodies. Confocal images showed decondensed extracellular chromatin at 8 hpi colocalizing with laminin and fibronectin within the fracture (Fig 3H, I). Collectively, this demonstrates that neutrophils responding to fractures release NETs, which interact with matrix proteins within the injury gap (Fig 3J).

To test whether neutrophils could play an active role in matrix management, comparable to that observed during soft tissue wound healing^10^, we performed immunohistochemistry for fibronectin, which is a major component of provisional matrix acting as a scaffold for later collagen deposition^32^. Confocal imaging showed that most neutrophils recruited to fractures contained fibronectin and laminin at 8 hpi (Figure 4A-D). 3D renders of neutrophils infiltrating fractures showed fibronectin and laminin spanning the cell membrane (Figure 4B). To further test the localisation of ECM proteins, we used a modular image analysis (MIA) pipeline ^33^, to test spatial distribution relative to the cell surface (Figure 4E, F). Mean fluorescence intensity of fibronectin and laminin showed increased levels of both matrix proteins within the cell at 4 and 8 hpi, compared to uninjured rays (Figure 4G, H), Further, levels of fibronectin, and laminin, were detected both at and up to 1um beyond the cell surface at 4hpi while only fibronectin spanned the membrane at 8hpi (Figure 4I, J), this shows that different matrix proteins have different spatiotemporal locations fibronectin frequently seen spanning the cell membrane, suggestive of it being moved or secreted.

**Figure 4.**
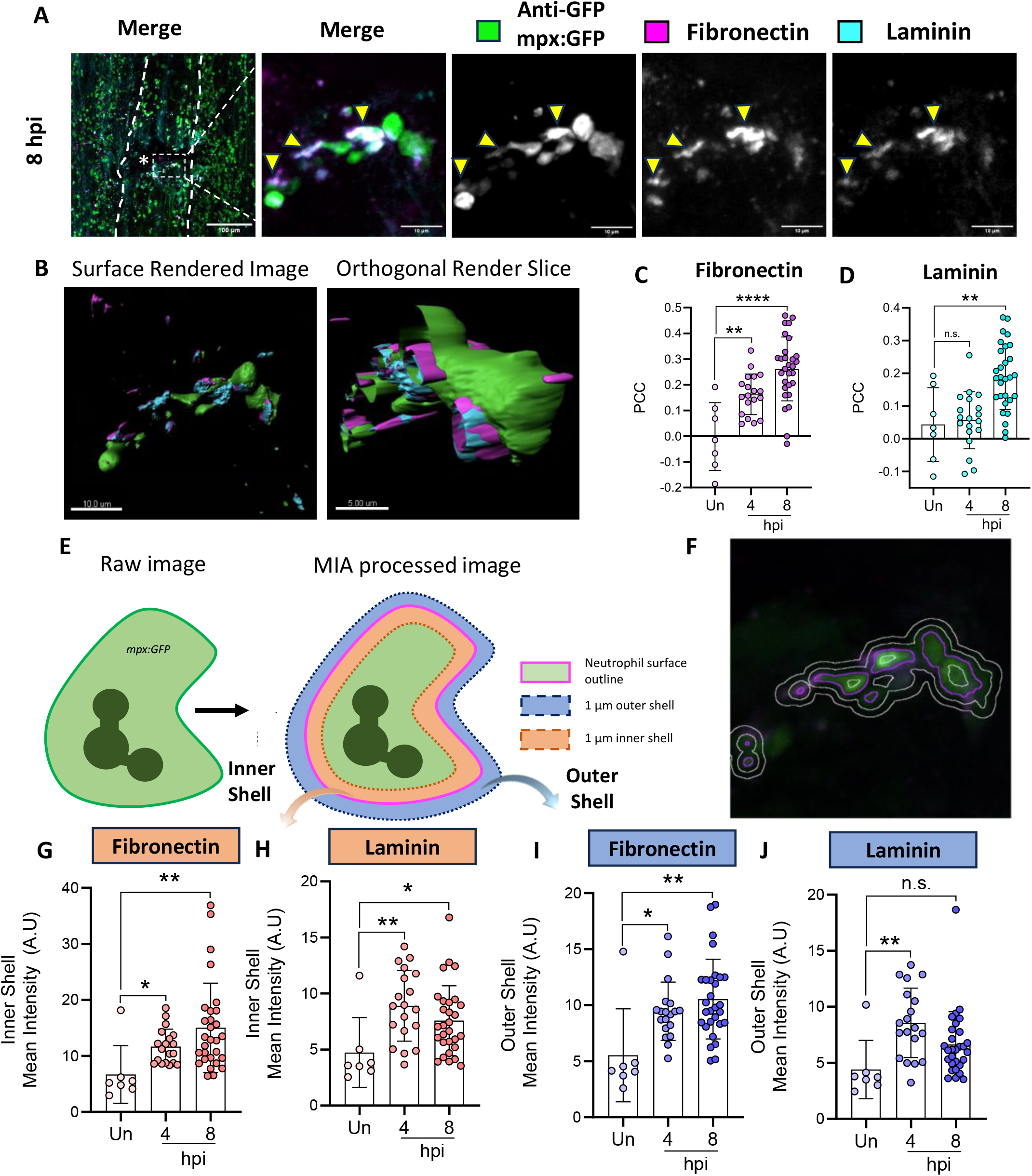
Neutrophils are rapidly recruited to fractures and stain positive for fibronectin and laminin. **A:** Representative confocal image of a fracture at 8 hpi immunostained for GFP (neutrophils) and the extracellular matrix components, fibronectin and laminin. Neutrophils recruited to fractures stain positive for fibronectin and laminin. Left scale bar = 100 um, right scale bar = 10 um. **B:** Imaris surface rendering from A shows ECM proteins above and below neutrophil surface. Left scale bar = 10 um, right scale bar = 5 um. **C-D:** Pearson’s correlation coefficient (PCC) was calculated to determine colocalization between GFP (neutrophils) and either fibronectin (B) or laminin (C). PCC values for both ECM proteins at 4 and 8 hpi, compared to uninjured fin tissue (Un). **E:** Schematic illustrating modular image analysis (MIA) method. **F:** Z slice of image from (A) after MIA processing; magenta = GFP segmentation, white rings = 1 μm and 2 μm shells, above and below cell surface. **G-J:** Results from MIA analysis showing mean intensity of either fibronectin (G & I) or laminin (H & J) in outer and inner 1 μm shells surrounding segmented neutrophils, which increases in all cases post fracture. Statistical analysis one way ANOVA comparing injured to uninjured, * = P < 0.05, ** = P < 0.01, n.s. = not significant.

To determine how neutrophils might be processing ECM proteins at the fracture site, we performed colocalization assays labelling with Rab7a to label late endosome/endolysosomal compartments in fracture tissue at 8 hpi. Both Rab7a and fibronectin showed higher abundance at the fracture site compared to uninjured distal tissue (Fig 5A-C). The percentage of neutrophils internalizing fibronectin within Rab7a vesicles, significantly increased following injury (Fig 5B-C), suggesting active internalisation for either degradation or recycling.

**Figure 5.**
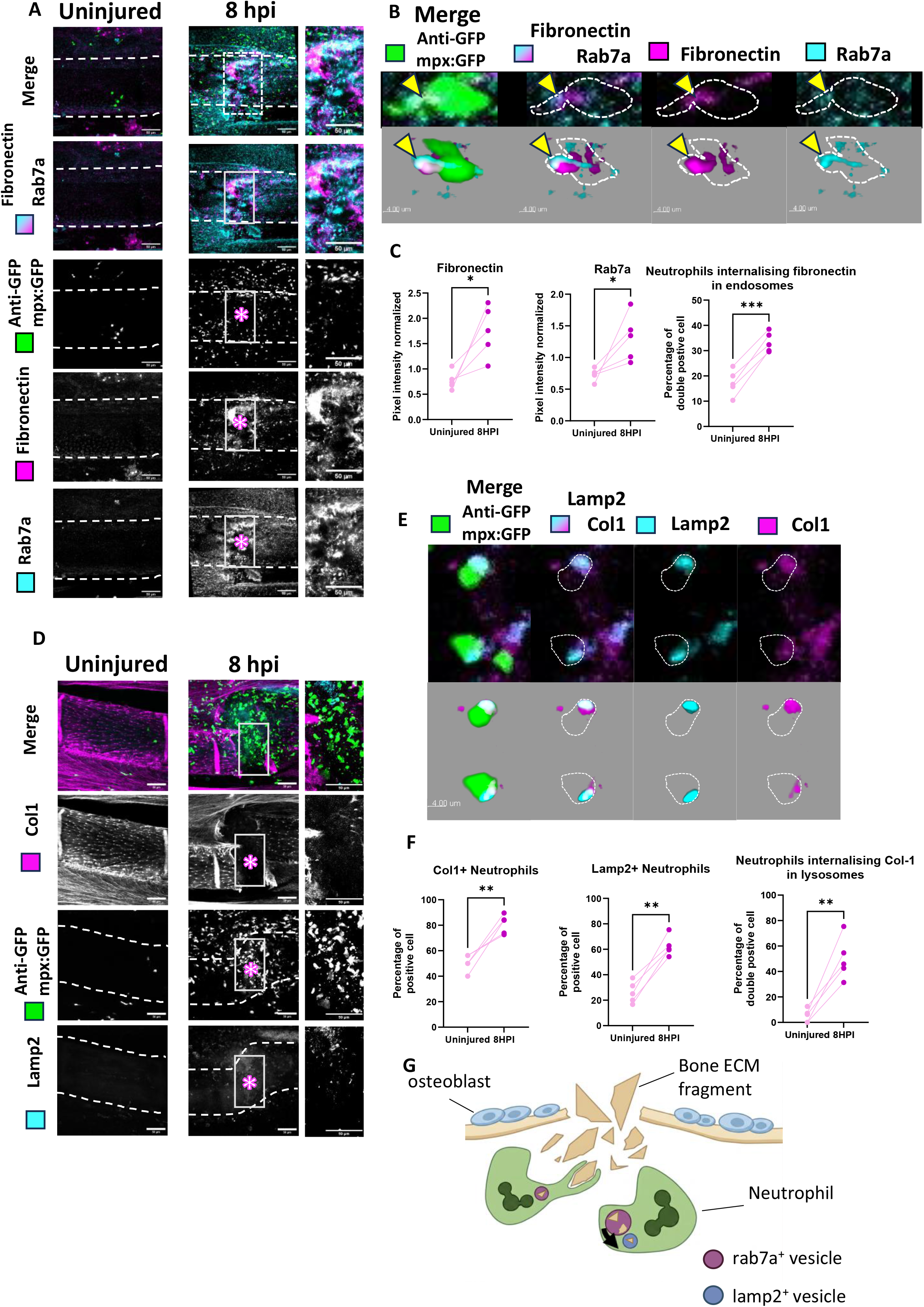
Neutrophils internalize ECM proteins within endosomes and lysosomes to organise the microenvironment at the fracture site. **A:** Representative confocal image of uninjured bone and fracture at 8 hpi immunostained for GFP (neutrophils), fibronectin and the endosome marker rab7a. Neutrophils were counted within a ROI (frame box) from the fracture site, of 80 μm width, 150 μm length and z of 45 μm. Scale bar = 50 μm. **B:** Single z focal plan and volume rendering, performed with Imaris, showing the internalization of fibronectin within rab7a vesicles, scale bar = 4 μm. **C:** The normalized mean pixel intensity for fibronectin and rab7a was measured in the uninjured bone and the ROI displayed in (A) of the injured bone of the same individuals. Neutrophils internalizing fibronectin within rab7a compartments during injury were manually counted within the ROI. Statistical analysis was performed using a Paired t-test, n=5, * = P < 0.05, ** = P < 0.01, *** =P < 0.001. **D:** Representative confocal image of uninjured bone and fracture at 8 hpi immunostained for GFP (neutrophils),), col1a1a and the lysosome marker lamp2. Neutrophils were counted within a ROI (frame box) from the fracture site, of 80 μm width, 150 μm length and z of 30 μm. Scale bar = 50 μm. **E:** Single z focal plan and volume rendering, performed with Imaris, showing the internalization of type I collagen within lamp2 vesicles. Scale bar = 4 μm. **F:** The number of col1a1a^+^ and lamp2^+^ neutrophils were manually counted in the uninjured bone and the ROI displayed in (D) of the injured bone of the same individuals. Neutrophils internalizing type I collagen within lamp2 compartments were manually counted within the ROI. Statistical analysis was performed using a Paired t-test, n=5, * = P < 0.05, ** = P < 0.01, *** =P < 0.001. **G.** Illustration representing neutrophils engaging in the reorganisation and internalisation of ECM fragments from the injury within rab7a^+^ and lamp2^+^ vesicles.

We performed further colocalization assays for Col1a1a and Lamp2. Lamp2 marks, endolysomal compartments but can also label secretory vesicles in neutrophils. We observed a dramatic increase both of Lamp2^+^ neutrophils and Col1a1^+^ neutrophils at the fracture site compared to uninjured tissue (Fig 5D, F). Injury also strikingly increased colocalization of Col1a1 with Lamp2^+^ neutrophil compartments from 5.179% to 49.98% (Fig 5E, F), consistent with enhanced trafficking of collagen through late endosomal and/or secretory compartments in activated neutrophils. Together these results are consistent with neutrophils playing an active role in matrix handling at the injury site (Fig 5G).

Neutrophils have recently been shown to play a role in matrix synthesis in the skin, helping to maintain barrier integrity ^34^. Lamp2 labels both endocytic and secretory vesicles in neutrophils, therefore, to test whether Col1a1 was being internalised or secreted, we examined neutrophil/collagen colocalization at different stages of neutrophil influx. Electron micrographs at the injury site, identified neutrophils containing fibrillar proteins with the periodicity of collagen, suggestive of internalisation or movement of mature collagen fibrils (Supplemental Fig 9A-C).

We also wanted to test whether neutrophils could be synthesizing Collagen, therefore we used the *col1a1a:GFP* promoter transgenic, which expresses GFP upon transcription of *col1a1a*, and observed expression in *lyz+* neutrophils (Supplementary Fig 10A). We also observed colocalization of Col1a1 with Golgi markers in neutrophils (Supplemental Fig 10B, C). Taken together these data establish that neutrophils can produce Type I Collagen at the site of injury, but interestingly they also appear to move mature Collagen fibrils.

## Discussion

Although the clinical literature remains conflicted, the weight of mechanistic and preclinical evidence indicates that NSAID exposure can adversely affect fracture healing ^35^. Rodent studies have shown selective COX-2 inhibition (coxibs) tends to show a stronger association with delayed healing/non-union than non-selective NSAIDs, with more pronounced effects observed with longer treatment duration ^36,37^. Human observational studies have reported increased non-union rates in NSAID-treated patients, particularly among users of COX-2 inhibitors, supporting concerns that suppression of the early inflammatory response may compromise successful bone regeneration ^38,39^. This raises the need for pre-clinical research to establish guidelines around safer practice on NSAID administration post-fracture. NSAIDs are globally overprescribed, accounting for ∼90% of all prescriptions for individuals over 65, a vulnerable population, with heightened fracture risk ^40^.

Here we demonstrate that neutrophils play a key role as architects of the injury microenvironment, interacting with matrix in multiple ways and undergoing NETosis at the site of injury. We propose that these functions collectively form an emergency scaffold, effectively templating repair until professional secretory cells, including stromal fibroblasts and osteoblasts can lay down a more permanent matrix. We build on previous studies of Cox2 inhibition to show that limiting neutrophil arrival at the site of injury, either pharmacologically or via genetic means, results in increased incidence of non-union fracture. Moreover, when union does occur, the resultant callus has abnormal architecture. This is expected to compromise mechanical performance of the repaired bone, in line with previous preclinical observations where NSAID administration at the time of fracture reduced mechanical strength at 21 days post fracture ^36^.

It is likely that the detrimental effect of NSAID use relative to injury is highly time dependent. Neutrophils are a transient population at the injury site; in our model, neutrophil numbers at the injury site decline by 24 hpi, suggesting that later modulation of neutrophils after the initial injury window would have a limited effect on bone outcomes. Indeed, prolonged neutrophil presence and sustained presence of NETs can be detrimental to outcomes ^27^. However, crosstalk with other immune cells and secretory cells in the stroma and with osteoblasts likely propagates early effects into the callus. Neutrophils have also been shown to directly modulate osteoblasts *in vitro* ^41,42^. Macrophages have been shown to instruct secretory cells in wounds, promoting differentiation and activation of fibroblasts, modifying matrix crosslinking ^43,44^, and to synthesise collagen themselves in certain wound contexts ^45^. We propose that the matrix synthesised by neutrophils is likely to be later modified both by macrophages and by the secretory populations that follow, and that this remodelling is beneficial for fracture repair.

Neutrophils are increasing viewed as a functionally heterogenous population ^46,47^. Our data supports this concept; we show that neutrophils play multiple roles at the fracture site. However, further investigation would be required to fully characterize the degree of heterogeneity. Here, we show that in fish, in line with human and mouse studies they can undergo NETosis at the fracture site ^27^. We further demonstrate that neutrophils act as matrix managers to modulate the local microenvironment, this aligns with rodent work showing that neutrophils can modulate matrix responses in skin and other soft tissue contexts ^34,48^. A wealth of research supports the concept of the ECM as an active player in repair and healing process orchestrating growth factor signalling, inflammatory cell recruitment and cell fate decisions rather than merely serving as a structural scaffold^49,50^.

Currently we cannot weight the relative functional importance of each neutrophil behaviour. However, by modulating the early fracture matrix, we propose neutrophils effectively constrain the environment, migration and mineralization of the subsequent osteoblast populations such that, despite their temporally restricted role, they have a lasting influence on the callus architecture and eventual bone outcomes.

## Materials and Methods

### Zebrafish Lines and husbandry

Zebrafish were maintained at the University of Bristol’s Animal Scientific Unit (ASU) and cared for following standard zebrafish husbandry guidelines (269). University of Bristol’s local Animal Welfare and Ethical Review Body (AWERB) and experiments conducted under a UK Home office Project License. Transgenic lines used are described in Table 1. Adult zebrafish used in this study were > 3 months post fertilization (mpf), < 12 mpf.

**Table 1.** Transgenic zebrafish lines used in this study, along with their origin.

| Line Name and Reference | Abbreviation | Description |
| --- | --- | --- |
| *Tg(mpx:GFP)<sup>i114</sup>* <sup>51</sup> | *mpx:GFP* | Neutrophils (cytoplasmic) |
| *Tg(lyz:h2a-mCherry)<sup>sh530</sup>* <sup>52</sup> | *lyz:h2a-mCherry* | Neutrophil chromatin (tagged) |
| *Tg(lyz:NTR-mCherry;mpx:GFP)<sup>i114</sup>*<br><sup>51,53</sup> | *lyz:NTR-mCherry;*<br>*mpx:GFP* | Neutrophils (cytoplasmic) – Targeted ablation |
| *Tg(Ola. Sp7:nlsGFP)<sup>zf13</sup>* <sup>54</sup> | *SP7:GFP* | Osteoblasts (Nuclear) |
| *Tg(lyz:NTR-mCherry; Ola. Sp7:nlsGFP)* <sup>53,55</sup> | *lyz:NTR-mCherry;*<br>*SP7:GFP* | Neutrophils (cytoplasmic) – Targeted ablation;<br>Osteoblasts (Nuclear) |
| *Tg(Col1a1a:GFP)* <sup>56</sup> | *col1a1a:GFP* | Collagen 1 expressing cells (cytoplasmic) |

### Anaesthesia

Tricaine methanesulfonate (MS222) stock solution was used for all anaesthesia. Stock solution was comprised of 12 mM Tris-HCl solution (SD8146; Bio Basic Canada) and 4 g L^−1^ MS222 powder (Sigma Aldrich) dissolved in Milli-Q water (Sigma Aldrich) pH adjusted to 7.5. Recovery anaesthesia was performed using MS222 stock solution dissolved in Danieau’s solution or aquarium water, at a final concentration of 160 mg L^−1^ for adults.

### Ibuprofen Treatment

Ibuprofen powder (I4883; Sigma Aldrich) was diluted in DMSO (Sigma Aldrich) to form a stock solution at a concentration of 100 mM made freshly on the morning of the first day of treatment. Immediately prior to treatment, stock solution was diluted into aquarium system water at final working concentrations. Adult zebrafish were treated with 10 µM via immersion from 24 hours prior to injury until the experiment end point or maximum 24 hours post fracture. Control zebrafish were immersed in system water containing an equivalent volume of DMSO. During treatment, 3 to 4 fish maximum were placed in a volume of 2L. Treatments were refreshed daily, and fish fed a standard diet.

### Cxcr2 Inhibition (SB225002)

SB225002 powder (Tocris, Bio-techne) was diluted in DMSO (Sigma Aldrich) to form a stock solution at a concentration of 100 mM which was stored in 100 µl aliquots at −20 °C for up to 1 year. Immediately prior to treatment, stock solution was diluted into aquarium system water at final working concentrations. Adult zebrafish were treated with 5 µM SB225002 via immersion from 24 hours prior to injury until the experiment end point. Control zebrafish were immersed in system water containing an equivalent volume of DMSO. Treatments were refreshed daily, and fish fed a standard diet.

### Neutrophil Ablation (Nifurpirinol)

Nifurpirinol (NFP) powder (17989855, Dr. Ehrenstorfer, Fisher Scientific) was diluted in DMSO (Sigma Aldrich) to form a stock solution at a concentration of 50 mM which was stored in 100 µl aliquots at −20 °C. Immediately prior to treatment, stock solution was diluted into aquarium system water at final working concentrations. Zebrafish were treated overnight with 2.5 μM NFP via immersion for 2 consecutive days (32 hours) prior to injury. Zebrafish were treated individually in a minimal volume of 150mL. Control zebrafish were immersed in system water containing an equivalent volume of DMSO. Treatments were refreshed each day, and each morning fish were placed back in standard housing conditions and fed a standard diet.

### Injury Models

To induce caudal fin fractures, adult zebrafish were anaesthetised in MS222 and placed laterally on a plastic petri dish under a brightfield stereomicroscope. The caudal fin was gently spread using a Pasteur pipette. A tweezer was used to crush a single lepidotrichium within the caudal fin causing fracture prior to the first bifurcation in the lepidotrichia and avoiding the 3 first dorsal and ventral rays of the fin. Where fin tissue was harvested for immunohistochemistry or proteomics, 3 to 4 fractures were induced in a single zebrafish, across the dorsoventral axis of the fin. Where repeated live time-course imaging was performed, 2 fractures were induced in a single zebrafish, across the dorsoventral axis of the fin. Every fracture was performed during the morning only.

### RNA collection and isolation

For RNA collection, 3 fractures were performed on 3 alternate hemi ray from the dorsal half of the caudal fin, and 3 fish were pooled together. Uninjured control coming from same individual were collected from the ventral half of the caudal fin having no fracture. The analysis was performed on triplicate for zebrafish having received no treatment, and on quadruplicate for treated fish (ibuprofen, SB225002, nifurpirinol). At 8hpi, caudal fins were amputated, resected into injured and uninjured, and collected in 400μl of ice-cold Trizol-LS (Thermofisher Scientific) at 8 hours post-injury (hpi). Fin samples were pulled through a 22G syringe () approximately 30 times to ensure proper maceration and stored at −80°C until RNA extraction. RNA was extracted via Trizol-Chloroform extraction and pellets were re-suspended in 20ul of UltraPure Distilled Water (Invitrogen). Ethanol precipitation was carried out overnight at −80°C to ensure RNA purity. RNA samples were centrifuged at 13000rpm for 30 minutes to pellet RNA and washed with pre-chilled 75% EtOH twice. Samples were suspended in a final volume of 15 ul of UltraPure water and measured with a nanodrop.

### RNA-seq analysis

Bulk RNA-seq analysis was performed tissue from control and fractured caudal fins (containing 3 fractures at 8 hpi) by Novogene. Sequencing reads were aligned to the *Danio rerio* reference genome (GRCz11, Ensembl release 109) using the corresponding Ensembl gene annotation ^57^.

Alignment and mapping were performed using the Novogene workflow. HISAT2 software was used to build the index of the reference genome and to align paired-end reads to it. The feature mapper used was featureCounts (2.0.6)^58^ to count read numbers mapped to each gene. Differential expression analysis was performed using DESeq2 R package (1.42.0)^59^. P-values were corrected for multiple testing using the Benjamini-Hochberg method. Differentially expressed genes (DEGs) were identified based on a log2FC and adjusted p-value ≤ 0.05.

### Gene enrichment analysis

Enrichment analysis was performed to assess whether DEGs identified in the above section were over-represented within cell-type-specific marker gene lists from a previously published zebrafish burn-injury study by Hou and colleagues, where they generated cell-type-specific marker gene lists based on scRNA-seq from FACS-sorted myeloid cells isolated from wounded and unwounded *Tg(mpx, mpeg1.1)* zebrafish larvae ^19^. The gene lists comprised markers for 10 myeloid cell clusters which were filtered to retain genes with a log2FC > 0.25 prior to enrichment analyses. The test sets consisted of 69 upregulated and 76 downregulated genes (injured ibuprofen vs dmso) and 46 upregulated and 21 downregulated genes (uninjured ibuprofen vs dmso) identified in the previous section.The background set comprised 19,988 unique zebrafish protein-coding genes. Upregulated and downregulated gene sets were analysed separately using hypergeometric tests, implemented using the testSeuratClusterEnrichment function in gRaphiaExtra R package (v0.26.8) deposited in GitHub: https://github.com/mskgrg/gRaphiaExtra^60^. Myeloid cell clusters were considered to be enriched if the test p-values passed the Bonferroni-adjusted threshold, correcting for the number of clusters tested ^60^ The analysis was repeated using all the gene symbols identified in the scRNA-seq analysis as the background population, yielding concordant results.

### Proteomics

#### Sample Collection for Proteomic

For proteomic analysis, 3 fractures were performed on the dorsal half of the caudal fin, and 3 fish were pooled together. Uninjured control coming from same individual were collected from the ventral half of the caudal fin having no fracture. Zebrafish within “Young uninjured” groups were not fractured and served as an additional control to test for potential systemic effects of fracture at distal sites in internal “control” caudal fin tissue. At 8hpi, caudal fins were amputated, resected into injured and uninjured if necessary, and placed into 200 μl of pre-prepared ice-cold protein lysis buffer with 10% Protease Inhibitor Cocktail (Roche, 04693116001). Samples were macerated using ultrasonication at 4°C and supernatants collected into fresh Eppendorf tubes, snap frozen in liquid nitrogen and stored at −20°C. Protein quantification was performed using Pierce BCA assay according to kit instructions and measured via nanodrop (Thermo Fisher, 23225). 15-Plex Tandem Mass Tagging proteomics were performed (Thermo Fisher).

#### TMT Labelling and High pH reversed-phase chromatography

Aliquots of 50µg of each sample were digested with trypsin, labelled with Tandem Mass Tag (TMTpro) sixteen plex reagents according to the manufacturer’s protocol (Thermo Fisher Scientific, Loughborough, LE11 5RG, UK) and the labelled samples pooled. An aliquot of 200ug of the pooled sample was desalted prior to fractionation by high pH reversed-phase chromatography using an Ultimate 3000 liquid chromatography system (Thermo Fisher Scientific). The sample was loaded onto an XBridge BEH C18 Column and the resulting fractions were analysed by nano-LC MSMS using an Orbitrap Fusion Lumos mass spectrometer (Thermo Scientific).

#### Nano-LC Mass Spectrometry

High pH RP fractions were fractionated using an Ultimate 3000 nano-LC system in line with an Orbitrap Fusion Lumos mass spectrometer (Thermo Scientific). Peptides in 1% (vol/vol) formic acid were injected onto an Acclaim PepMap C18 nano-trap column (Thermo Scientific). After washing with 0.5% (vol/vol) acetonitrile 0.1% (vol/vol) formic acid peptides were resolved on a 500 mm × 75 μm Acclaim PepMap C18 reverse phase analytical column (Thermo Scientific). All spectra were acquired using an Orbitrap Fusion Lumos mass spectrometer controlled by Xcalibur 3.0 software (Thermo Scientific) and operated in data-dependent acquisition mode using an SPS-MS3 workflow. FTMS1 spectra were collected at a resolution of 120 000, with an automatic gain control (AGC) target of 200 000 and a max injection time of 50ms. The MS2 precursors were isolated with a quadrupole isolation window of 0.7m/z. ITMS2 spectra were collected with an AGC target of 10 000, max injection time of 70ms and CID collision energy of 35%. For FTMS3 analysis, the Orbitrap was operated at 50 000 resolution with an AGC target of 50 000 and a max injection time of 105ms. Precursors were fragmented by high energy collision dissociation (HCD) at a normalised collision energy of 60% to ensure maximal TMT reporter ion yield. Synchronous Precursor Selection (SPS) was enabled to include up to 10 MS2 fragment ions in the FTMS3 scan.

#### Proteomic Analysis

The raw data files were processed and quantified using Proteome Discoverer software v2.4 (Thermo Scientific) and searched against the UniProt Danio rerio database (downloaded January 2022: 61622 entries) using the SEQUEST HT algorithm. Peptide precursor mass tolerance was set at 10ppm, and MS/MS tolerance was set at 0.6Da. Searches were performed with full tryptic digestion and a maximum of 2 missed cleavages were allowed. The reverse database search option was enabled, and all data was filtered to satisfy false discovery rate (FDR) of 5%. Protein groupings were determined by PD2.4, however, protein annotation was improved using an in-house script which reselects the master protein for each group based upon the current uniprot annotation score of the candidate master proteins.

#### Whole mount Immunohistochemistry

Tissues were fixed in 4% paraformaldehyde (PFA) for 2 hours at room temperature or overnight at 4°C and then dehydrated using a serial dilution of methanol (MeOH) in PBS-Tx (0.2% Triton X in Phosphate Buffered Saline). Samples were left in 100% MeOH for 10 minutes at room temperature prior to rehydration or stored at −20°C in 100% MeOH long term. Tissues were rehydrated in a serial dilution of MeOH in PBS-Tx. After washing 3 times, samples were digested with Proteinase K ([1:1000]; P5568; Sigma-Aldrich) diluted in PBS-Tx for 90-120 minutes at room temperature. Proteinase K solution was refreshed every 30 minutes. Samples were washed 3 times in PBS-Tx and then incubated in blocking buffer at room temperature for 3 hours (5% horse serum in PBS-Tx). Samples were then incubated with primary antibodies overnight at 4°C with gentle agitation. Samples were washed 6 times in PBS-Tx before being incubated in blocking buffer for 2 hours and then stained with secondary antibodies at room temperature for 2 hours. All antibodies were diluted in blocking buffer. Primary antibodies used were: chick mAb to GFP ([1:500]; ab13970; Abcam), rat mAb to mCherry ([1:100]; 16D7; Invitrogen), rabbit mAb to laminin ([1:200]; Abcam; ab11575), mouse mAb to fibronectin ([1:200]; Sigma-Aldrich; F7387); mouse mAb to NET-specific histone H3 cleavage ([1:200]; 3D9; Max Planck Institute, Berlin4242), mouse mAb to Lamp2 ([1:200]; H4B4; DSHB), rabbit pAb to Col1a1a ([1:200]; Genetex; GTX133063), rabbit pAb to myeloperoxidase ([1:200]; Genetex; 128379). Secondary antibodies used were: goat anti-chick Alexa Fluor488; goat anti-rat Alexa Fluor-555 goat anti-mouse; Alexa Fluor-555; goat anti-mouse Alexa Fluor-568; donkey anti-rabbit Alexa Fluor 647 (A11039; A21434; A21137; A21124; A31573, Thermo Fisher Scientific). Samples were then washed a final 6 times in PBS-Tx. For nuclear staining, samples were incubated in DAPI ([1:2000]; D1306; Thermo Fisher) and washed a further 3 times. Stained samples were stored in PBS-Tx at 4°C. All washes were performed for 10 minutes at room temperature, unless stated otherwise. Stained samples were imaged on a confocal microscope within 2 weeks of staining.

#### Stereomicroscope Imaging

Fish were immobilised via anaesthesia at the required time-points post-injury placed on a plastic petri dish. Images of the fin were taken in the dark using a DFC700T camera mounted to a MZ10F Stereomicroscope (Leica Microsystems). Images were acquired using LAS X software (version 3.7.0). Control images of uninjured fin tissue was obtained at each time point to normalise for variation in transgene reporter efficiency and image acquisition settings. Fish were recovered in fresh system water after imaging.

#### Confocal imaging

Fixed tissues were mounted in 0.3% UltraPure™ Low Melting Point Agarose (Thermo Fisher) dissolved in PBS. Samples were imaged with a x10 or x20 objective lens on a SP5 confocal microscope (Leica Microsystems). Images were acquired using LAS X software (version 3.0 or above). For 3-dimensional imaging, confocal stacks were taken between 1 to 2 µm increments through the tissue.

#### Spinning disk imaging

Fractures were performed on the morning, and caudal fin were resected approximatively 3 hours post-injury. The caudal fin was places in 0.5% UltraPure™ Low Melting Point Agarose (Thermo Fisher) dissolved in culture medium as described in ^62^ to allow *ex-vivo* imaging. Imaging was performed with the Yokogawa CSU-W1 SoRa spinning disk for 2 hours. Images were acquired every 1min20 second, with a z-slice of 1μm over 180 μm with dual cameras model T2 to capture simultaneously GFP and mCherry channels; and brightfield images separately.

#### High resolution μCT (nanoCT)

Samples were mounted between two adhesive Kapton (polyimide) sheets and held vertically in a keyless chuck holder. The holder was positioned inside a Zeiss Xradia 520 Versa µCT scanner operating at 7W and 80kV with no filtering. Preliminary scans were taken using 201 projections to verify positioning and minimise any movement from sample relaxation and/or heating effects from the source. The 20 × magnification lens was used with 2400 projections to achieve sub-500 nm voxel resolution.

#### Image Analysis

3D image rendering and animations were constructed using Imaris image analysis software. FIJI was used for all other analyses and measurements^63^. Intensity ratios were calculated by measuring the average pixel intensity of injured tissue within a region of interest (ROI), divided by average pixel intensity of uninjured tissue within an equivalent ROI taken using the same exposure settings, as previously described^16^. This method of quantification normalises for any variability in reporter expression between animals or image acquisition settings from day-to-day, allowing for more reliable comparison of fluorescence intensities between zebrafish. The ModularImageAnalysis (MIA) plugin was used for automated analysis of ECM proteins with respect to neutrophils ^64^. Neutrophils were segmented using a GFP confocal image stack, which was first processed with a 2D Gaussian filter to remove noise, then binarised with a user-defined intensity threshold. Holes and regions in the binarised image smaller than 5 µm^3^ were removed prior to the image being passed through a 2D median filter to smooth region edges. A pair of shells around the binarised regions were created; the first extending from the surface of the binarised regions up to 2 µm away and the second from 2 to 4 µm away^33^. Both the 3D shell volume and fluorescence intensity in the corresponding ECM signal channels was measured. Pearson’s correlation coefficient colocalization for two ECM signals was also measured both across the entire image stack and in masked regions, where only pixels with some GFP or ECM signal were considered.

### Statistical Analysis

GraphPad PRISM Software (version 11.1.0) was used for all statistical analyses and graph design. Normality was evaluated with a Shapiro-Wilk to determine whether a parametric or nonparametric statistical test should be used. To compare a single time-point, between internal uninjured control and bone injury, a Two-tailed Paired t-test (parametric) was used. Where more than two sets measurements were compared (e.g., different fish at multiple time points), a one-way ANOVA (parametric) or Kruskal-Wallis’s test (non-parametric) with Tukey’s correction was used. Where more than two sets measurements were compared for different groups (e.g., different fish at multiple time points and for different treatment), a Two-way ANOVA with Tukey’s correction was used. For comparison of frequencies between groups (e.g., frequency of fracture dissociation), a Chi-square analysis was performed. Where repeat measures were taken from the same animal over time, such as throughout a live time course of fracture repair, statistical tests were adjusted to account for repeated measures. Differences were considered statistically significant where P < 0.05, as per convention. Graphs display mean ± the standard deviation as error bars (where appropriate) or shows pairing of measure between internal uninjured control and their respective measure in bone injury.

## Supporting information

Supplemental Video 1A. Real-time in vivo video of neutrophils performing NETosis at the fracture site in Tg(mpx:GFP; lyzH2a:mCherry) fish.

Supplemental Video 1B. Volume rendering of a neutrophil performing NETosis at the fracture site, isolated from supplemental video 1A.

Supplemental Video 2. Video from electron micrograph with bone in cyan, enucleated neutrophil in magenta, and nucleated neutrophil in orange.

Supplemental Video 3. Neutrophils (green) perform NETosis as they extrude a 3D9 positive scaffold (magenta) colocalising with mpx granules (cyan).

## Funding and Acknowledgements

**JZ** and **BF** were funded by Vivensa AISRPG2305\7 awarded to **BA** and **CLH**. **LMM** and **RC** were funded by the Wellcome Trust Dynamic Molecular Cell Biology PhD Programme at the University of Bristol (No. 108907/Z/15/Z), **LMM** was subsequently funded by Dunhill Medical Trust Seed Funding Award (AIS2110\25) awarded to **BA** and **CLH. CLH** was funded by Versus Arthritis Senior Research Fellowship (29137). **NRD** is supported by an Australian Government Research Training Program Scholarship and the Mater Research Frank Claire Scholarship. **JPK** is funded by a National Health and Medical Research Council (Australia) Investigator Grant (GNT2026272) and the Mater Foundation. We gratefully acknowledge Dr Kaitlyn A. Flynn’s support. We thank Dr Bernadette Caroll and Dr Nicola Stevenson for providing antibodies against lamp2, rab7a and golgin-160. We acknowledge the Wolfson Bioimaging Facility for imaging support and the Bristol Proteomics facility for aiding proteomics experiments. We acknowledge the help of Dr Chris Neal for the electron micrographs acquisition.

## Contributions

**JZ:** Conceptualization; data curation; formal analysis; investigation; methodology; resources; validation; visualization; funding acquisition; writing-original draft; writing-review and editing. **LMM**: Conceptualization; data curation; formal analysis; investigation; methodology; resources; validation; visualization; funding acquisition; writing-original draft. **RMC:** Conceptualization; investigation, methodology. **RS**: investigation, methodology. **NRD**: investigation, methodology. **BHF:** investigation, methodology. **BTG**: formal analysis; methodology. **JJM**: investigation; methodology. **SC**: Formal analysis; methodology; software. **JPK**: investigation; methodology. **BA**: Conceptualization; data curation; funding acquisition; project administration; resources; supervision; writing-review and editing. **CLH**: Conceptualization; data curation; funding acquisition; project administration; resources; supervision; writing-review and editing.

## Data Availability

Imaging data and .mia MIA plugin workflow files are available through the University of Bristol’s RDSF server. Further information and requests for materials associated with this study should be directed to and will be made available upon reasonable request by the lead contact, Chrissy Hammond at. All raw data will be available via a DOI at data.bris.ac.uk upon article acceptance.

---

**Supplementary Figure 1.**
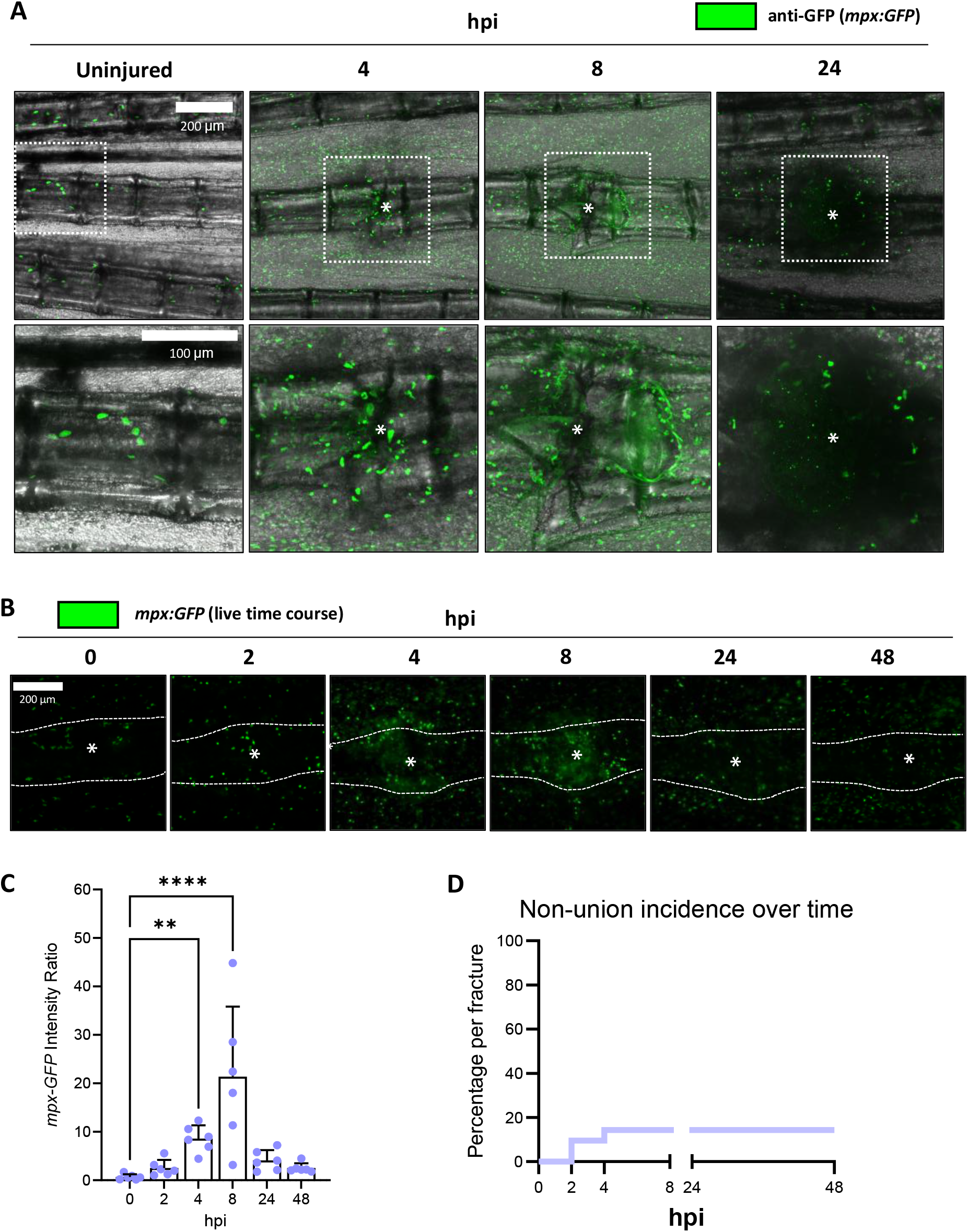
Time course of neutrophil response to fracture and the proportion of non-union. Fractures were induced in the caudal fins of adult *mpx:GFP* zebrafish. **A:** Representative confocal images of uninjured bone and fractures at 4, 8 or 24 hours post injury (hpi) immunostained for GFP (neutrophils). The (*) indicates the centre of the fracture. **B:** Representative stereomicroscope images of fractures in *mpx:GFP* transgenic zebrafish followed longitudinally between 0 and 48 hpi. The (*) indicates the centre of the fracture. Scale bar = 200 μm **C:** Neutrophil recruitment to fractures was measured via relative fluorescence intensity, whereby GFP intensity at the fracture site was measured relative to GFP intensity in uninjured bone within the same fish, n=6. Statistical analysis was performed using Kruskal-Wallis’s test comparing time points. **** = P < 0.0001, ** = P < 0.01. **D:** Occurrence of non-union fracture through the loss of the injured ray was reported at 2,4,8,24 and 48 hpi. The proportion of non-union over union was plotted per fracture, over 42 fractures in 21 fish (2 fractures/fish).

**Supplemental 2:**
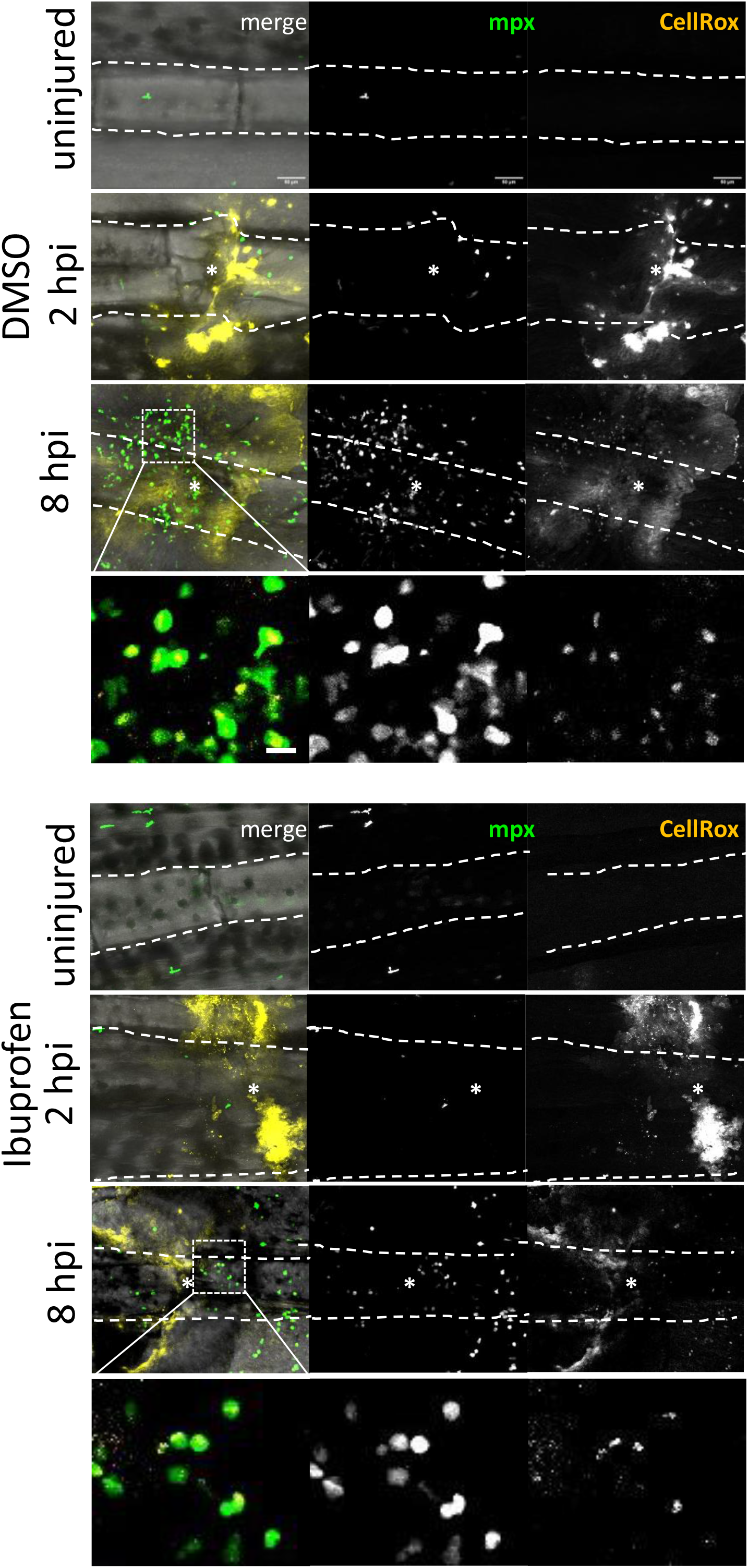
Ibuprofen treatment (10μM) does not prevent ROS production by neutrophils at the injury site. Evaluation of ROS production in *mpx:GFP* zebrafish treated with Ibuprofen compared to control. Confocal microscopy of the uninjured caudal ray (dash line) and the injured ray collected at 2 hpi and 8 hpi. Scale bar = 50 μm. Magnification on ROS-positive neutrophils at 8 hpi. Scale bar = 10 μm.

**Supplemental Figure 3.**
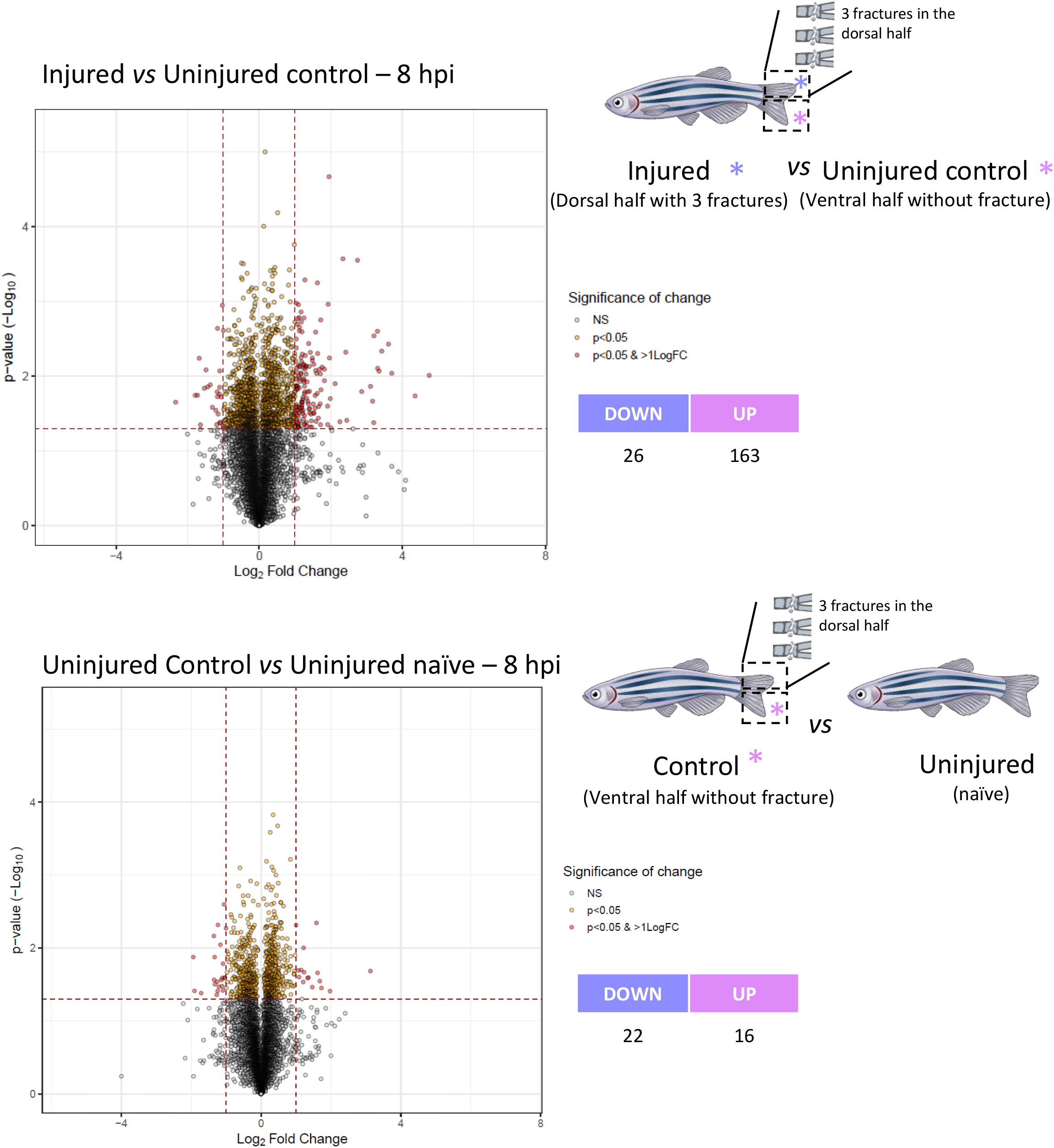
Evaluation of the systemic effects induced by fractures over uninjured area. Comparison of the uninjured ventral half of the caudal fin used as control in the proteomic analysis of bone fractures (Control group) versus zebrafish that never experienced caudal fin fracture (Uninjured group). Volcano plot of the significantly regulated proteins in the control compared to the uninjured fish (p-adj < = 0.05 and |log_2_foldchange|>= 1.

**Supplemental Figure 4.**
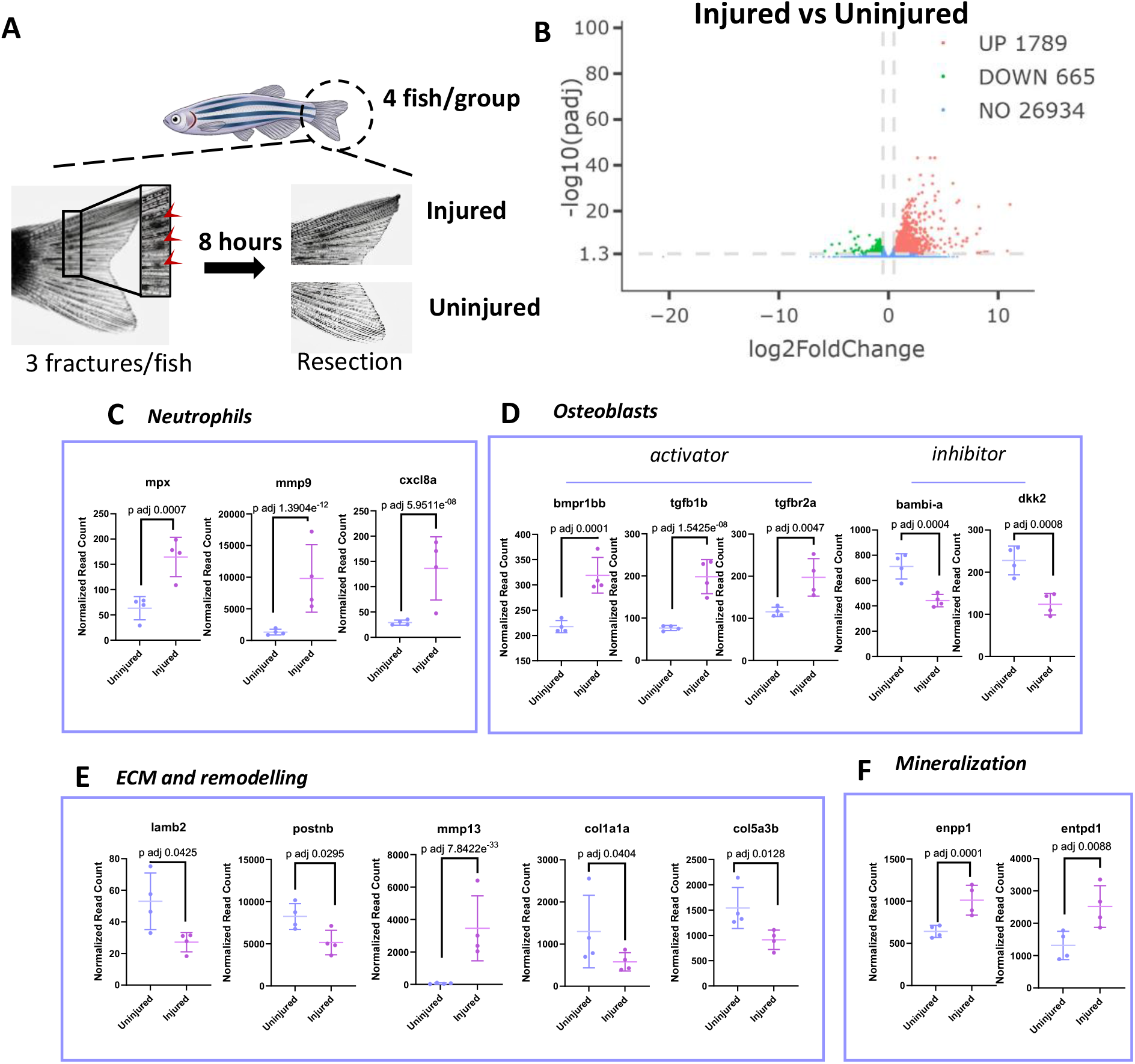
Evaluation of the early transcriptomic modifications following a bone injury at 8 hpi. **A.** Schematic representation of the sample collected for transcriptomic. **B:** Vulcano plot of the differentially expressed genes (DEG) in injured vs uninjured control (p-adj < = 0.05 and |log_2_foldchange|>= 0.5. **C:** Comparison of neutrophil related DEG in injured vs uninjured control. Normalized read count are plotted with SD. **D:** Comparison of osteoblast related DEG in injured vs uninjured control. Normalized read count are plotted with SD. **E:** Comparison of ECM and remodelling related DEG in injured vs uninjured control. Normalized read count are plotted with SD. **F:** Comparison of mineralization related DEG in injured vs uninjured control. Normalized read count are plotted with SD.

**Supplemental Figure 5.**
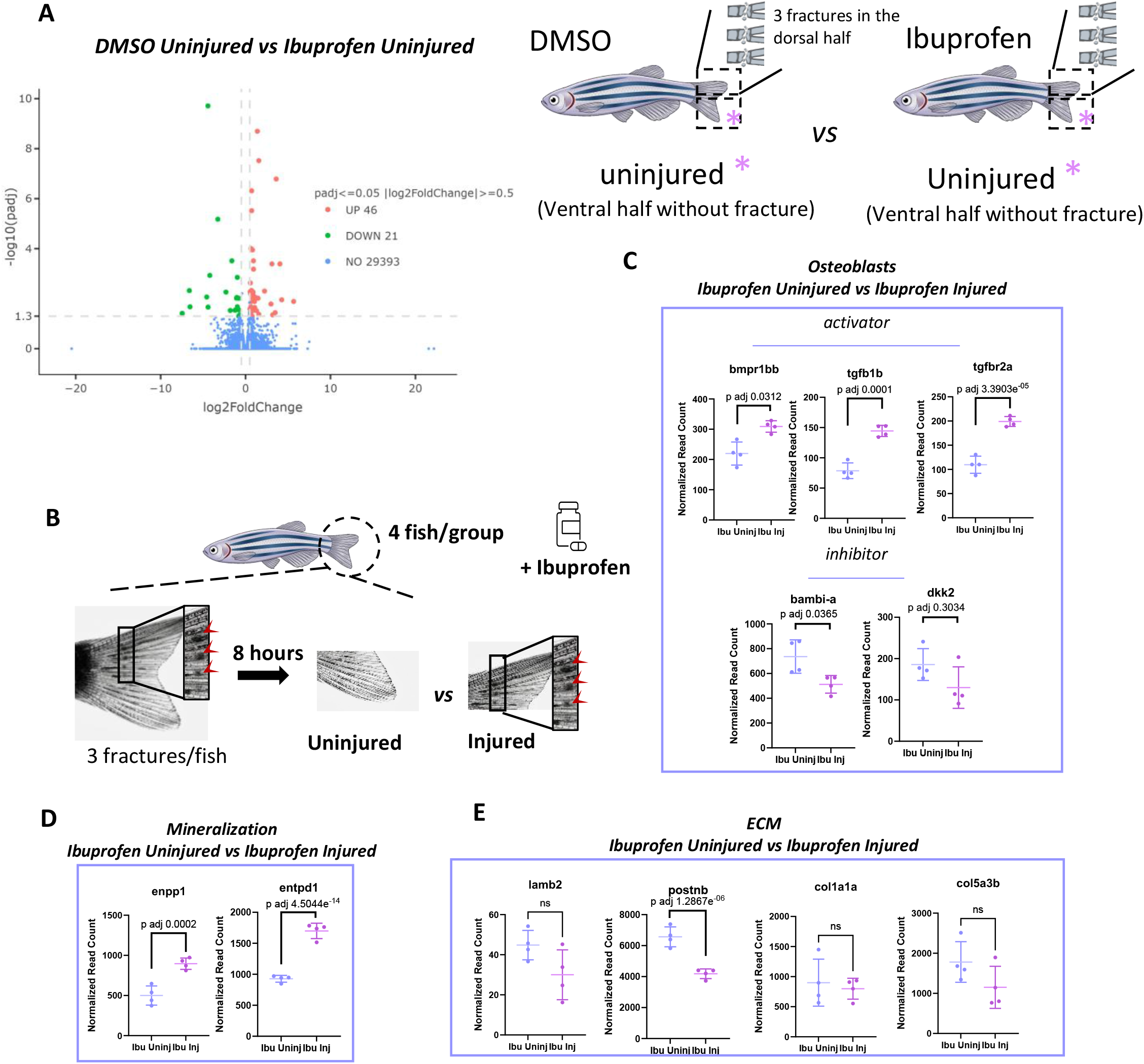
Evaluation of the systemic effects induced by ibuprofen over uninjured area and the effects of ibuprofen following bone injury. **A.** Vulcano plot of the uninjured area from ibuprofen treated zebrafish against the DMSO control at 8hpi (p-adj < = 0.05 and |log_2_foldchange|>= 0.5). **B:** Schematic representation of the samples collected for the transcriptomic comparison of ibuprofen uninjured vs Ibuprofen injured at 8 hpi. **C:** Comparison of osteoblast related DEG in ibuprofen injured vs ibuprofen uninjured control. Normalized read count are plotted with SD. **D:** Comparison of mineralization related DEG in ibuprofen injured vs ibuprofen uninjured control. Normalized read count are plotted with SD. **E:** Comparison of ECM related DEG in ibuprofen injured vs ibuprofen uninjured control. Normalized read count are plotted with SD.

**Supplemental Figure 6:**
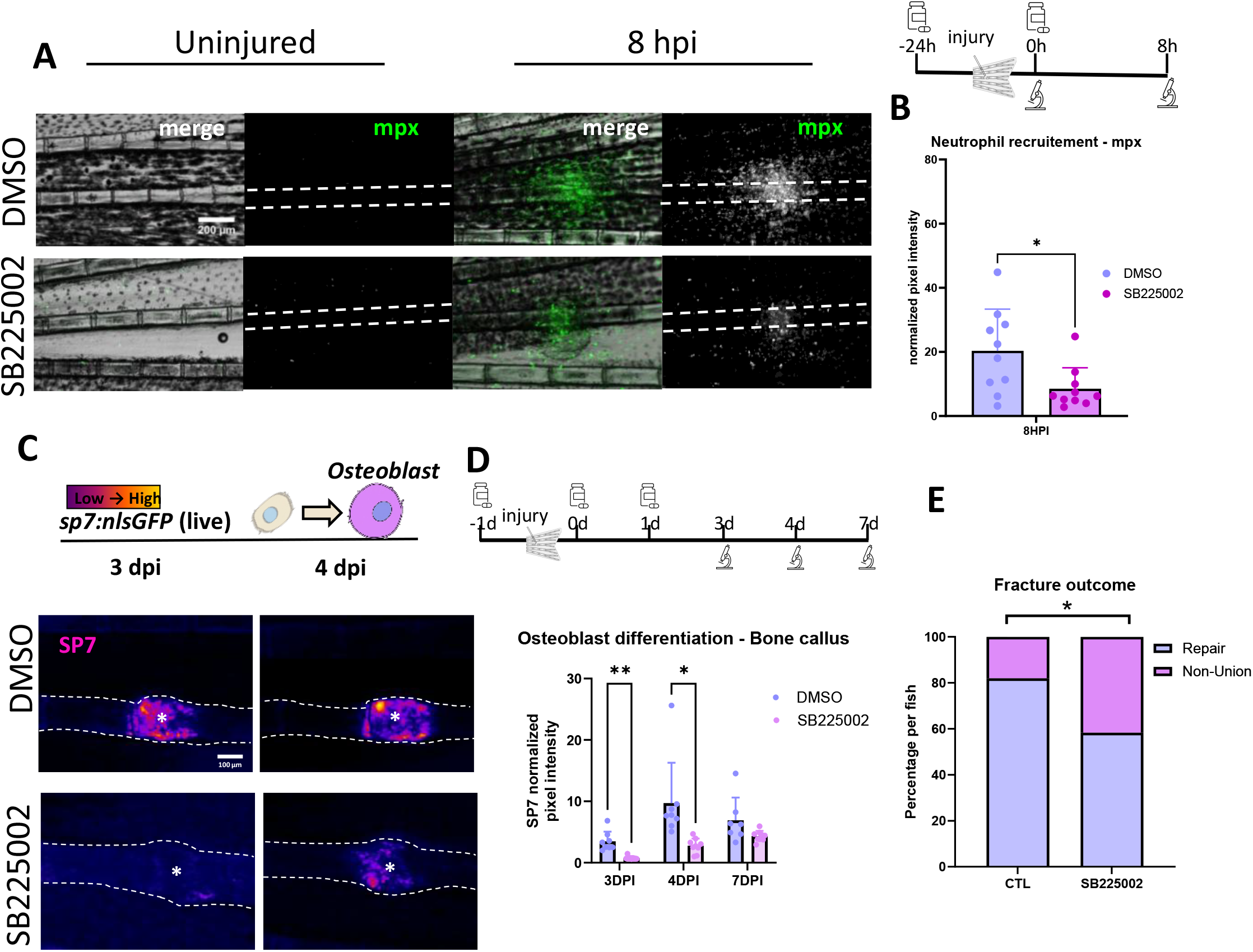
Blocking neutrophil recruitment with SB225002 treatment (5μM) recapitulates the phenotype of ibuprofen treated fish and negatively impacts the bone repair outcome. **A:** Representative stereomicroscope image of uninjured and injured bone at 8 hpi in *Tg(mpx:GFP)* zebrafish treated with SB225002 compared to DMSO control. Scale bar = 200 μm. **B:** The normalized pixel intensity for *mpx:GFP* is displayed as a bar plot with SD, n=10 per group. Statistical analysis was performed with a Mann Whitney test, * = P < 0.05. **C:** Representative stereomicroscope images of the intensity of expression of the osteoblast marker *sp7* at 3 and 4 dpi comparing DMSO control and SB225002 treated fish. The (*) marks the centre of the injury. Scale bar = 100 μm. **D:** Fish were longitudinally followed from 0 to 7 dpi for measuring *sp7* expression. The normalised *sp7* pixel intensity is displayed as a bar plot, n=7-9 per group. Statistical analysis was performed with a Tukey’s multiple comparisons test, ** = P < 0.01, * = P < 0.05. **E:** Percentage of the incidence of non-union fracture following the SB225002 treatment, per individual. Statistical analysis was performed with a Chi-square, n = 50 for the control group and n = 24 for the SB225002 group.

**Supplemental Figure 7:**
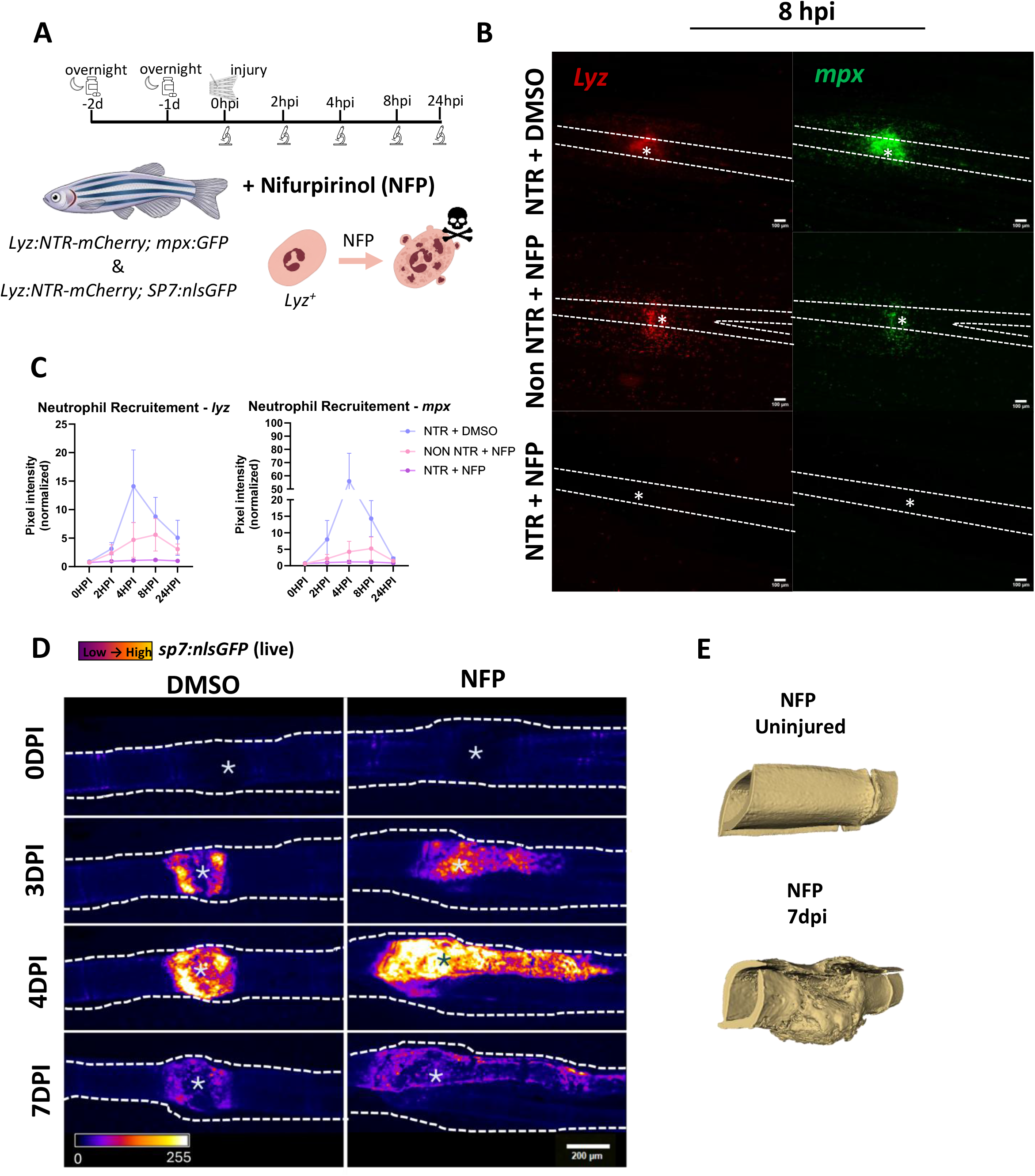
Genetic ablation of neutrophils leads to alteration of the bone repair mechanism. **A:** Specific *lyz^+^* neutrophil ablation can be induced in *Tg(lyz:NTR-mCherry)* by adding the nifurpirinol (NFP) pro-drug resulting in the death of cell expressing the nitroreductase (NTR) under the *lyz* promotor. Prior the injury, zebrafish are treated with NFP (2.5μM) during 2 overnights and then placed back in fresh water. **B:** Representative stereomicroscope image of uninjured and injured bone at 8 hpi in *Tg(lyz:NTR-mCherry; mpx:GFP)* treated with DMSO (NTR +DMSO) or NFP (NTR +NFP) ; and in *Tg(lyz:dsRed; mpx:GFP)* treated with NFP (non NTR +NFP) as a second control group against the pro-drug. The (*) marks the centre of the injury. Scale bar = 100 μm. **C:** Fish were longitudinally followed from 0 to 24 hpi for measuring *lyz* and *mpx* normalised pixel intensity, n=3 for DMSO, n=4 for non NTR + NFP, n=6 for NTR + NFP. **D:** Representative stereomicroscope images of the intensity of expression of the osteoblast marker *sp7* at 0, 3, 4 and 7 dpi in *Tg(lyz:NTR-mCherry; sp7:nlsGFP)* treated with DMSO or NFP. Fish treated with NFP showed bone repair alteration with disorganised callus. The (*) marks the centre of the injury. Scale bar = 200 μm. . **E:** Volume rendering of high resolution nanoCT of uninjured bone and bone calluses at 7 dpi.

**Supplemental Figure 8.**
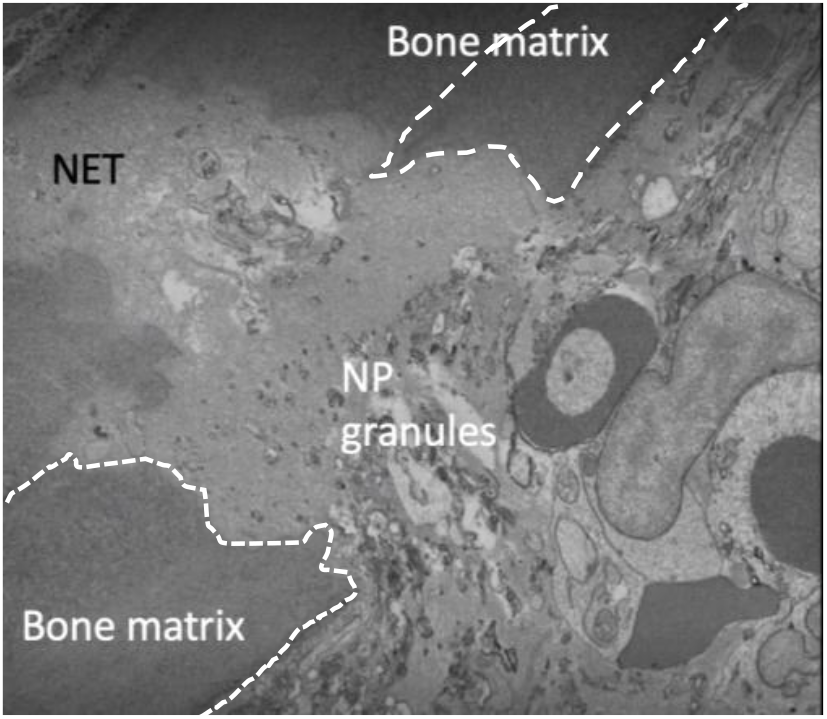
Enucleated neutrophils are located at the fracture site indicating they undergo NETosis. Representative EM image of a fracture at 8hpi. The bone matrix is outlined by the dotted line, neutrophils granules and NET are indicated on the image.

**Supplemental Figure 9.**
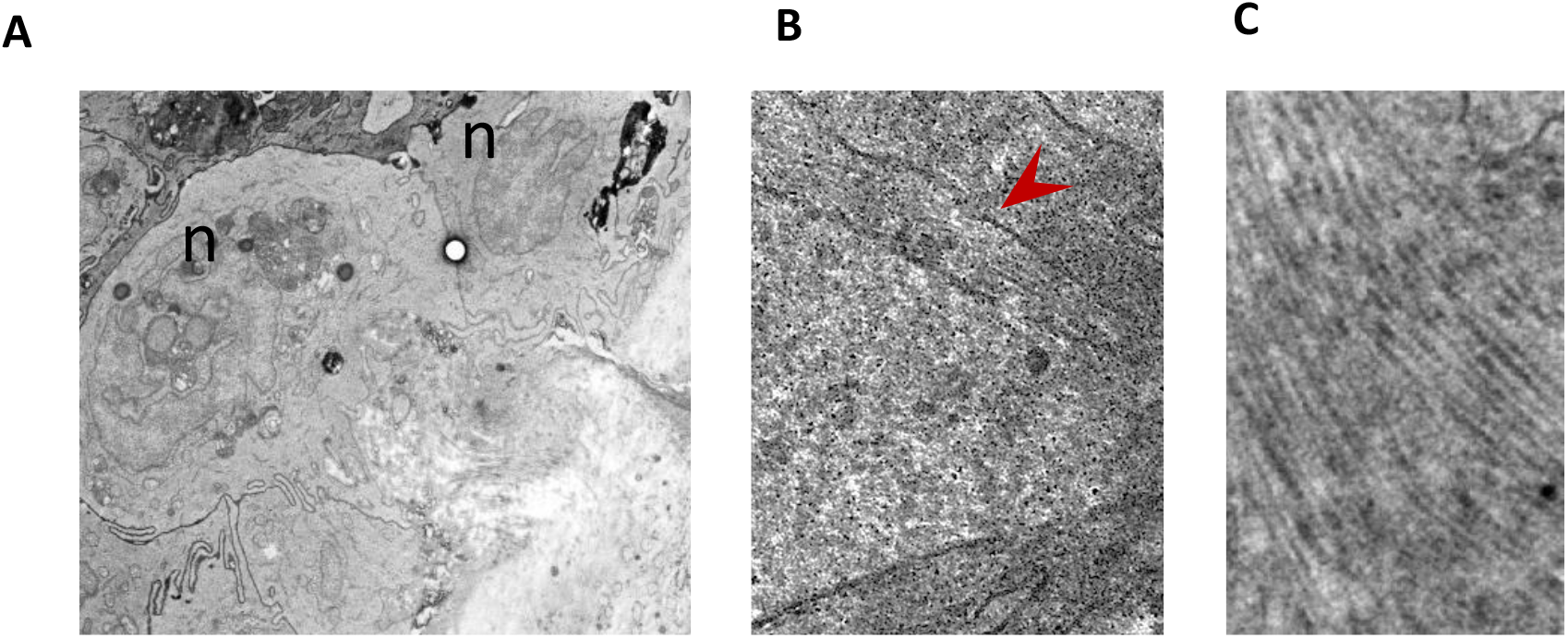
Neutrophils internalize collagen at the fracture site. **A.** Representative EM image of neutrophils at the fracture site. n=multi lobed nucleus. **B.** The red arrowhead indicates membrane-bounded compartment. **C.** Image showing internalization of collagen within a neutrophil preserving its periodic banding.

**Supplemental Figure 10:**
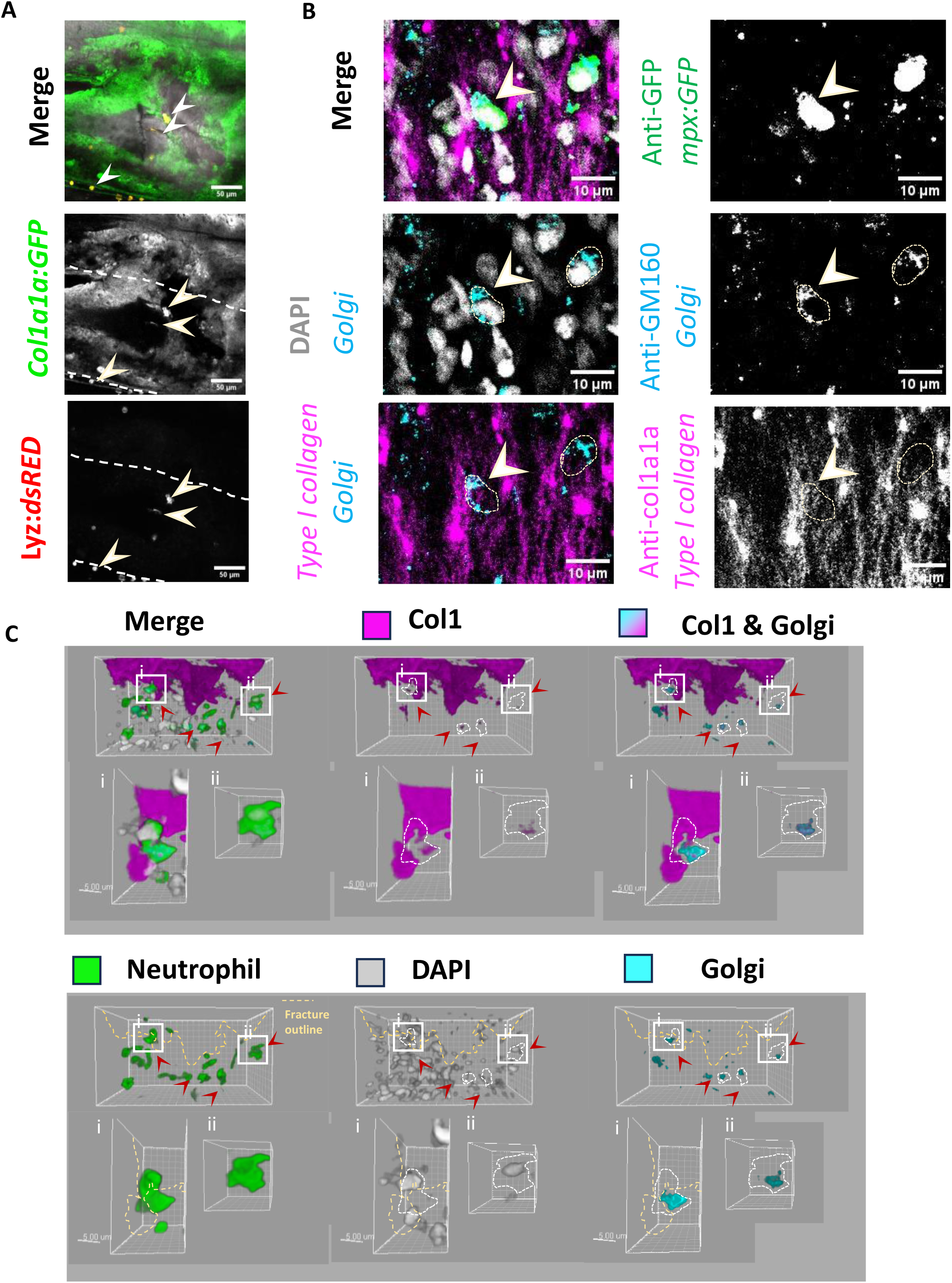
Neutrophils express type I collagen at the fracture site. **A.** Representative confocal image of a fracture at 8 hpi in *Tg*(*col1a1a:GFP; lyz:dsRed)* zebrafish. Lyz^+^ neutrophils expressed col1a1a and are located inside the fracture as indicated by the red arrowheads. Scale bar = 50 μm. **B.** Fracture at 8 hpi in *mpx:GFP* fish immunostained for GFP (neutrophils), GM160 (cis-Golgi), col1a1a and stained with DAPI (nucleus). The anti GM-160 antibody revealed large perinuclear golgi structure. The white arrowhead shows a neutrophil colocalising collagen within its Golgi apparatus, indicating for its synthesis. Scale bar = 10 μm. **C.** Volume rendering from (B). The yellow dotted line indicates the fracture outline. Red arrowheads show neutrophils synthetising type I collagen at the proximity of the fracture outline with (i) and (ii) focusing on specific cells showing colocalization of collagen within their Golgi apparatus. Scale bar = 5 μm.

**Supplemental Video 1. Real-time *in vivo* video of neutrophils performing NETosis at the fracture site. A:** Spinning disk video showing neutrophil recruitment toward the bone fracture in *Tg(mpx:GFP; lyzH2a:mCherry)* fish; with *mpx* labelled in cyan and *lyzH2a* in magenta. Neutrophils in proximity of the fracture site extruded their chromatin as evidence of NETs. Scale bar = 50 μm. **B:** Video of the volume rendering with Imaris of a neutrophil performing NET isolated from the spinning disk video (A).

**Supplemental video 2. Video from electron micrograph of a fracture at 8hpi.** Bone is labelled with cyan, enucleated neutrophil with magenta, and nucleated neutrophil with orange.

**Supplemental Video 3. Neutrophils perform NETosis as they extrude a 3D9 positive scaffold colocalising with Mpx granules.** Video of the volume rendering with Imaris of fracture at 8hpi of a *Tg(mpx:GFP)* fish immunostained for GFP labelling neutrophils (green), Mpx granules (cyan) and NETs with the 3D9 antibody (magenta). Scale bar = 50 μm

